# HuD (ELAVL4) gain-of-function impairs neuromuscular junctions and induces apoptosis in *in vitro* and *in vivo* models of amyotrophic lateral sclerosis

**DOI:** 10.1101/2023.08.22.554258

**Authors:** Beatrice Silvestri, Michela Mochi, Darilang Mawrie, Valeria de Turris, Alessio Colantoni, Beatrice Borhy, Margherita Medici, Eric Nathaniel Anderson, Maria Giovanna Garone, Christopher Patrick Zammerilla, Udai Bhan Pandey, Alessandro Rosa

## Abstract

Early defects at the neuromuscular junction (NMJ) are among the first hallmarks of the progressive neurodegenerative disease amyotrophic lateral sclerosis (ALS). According to the “dying back” hypothesis, disruption of the NMJ not only precedes, but is also a trigger for the subsequent degeneration of the motoneuron in both sporadic and familial ALS, including ALS caused by the severe *FUS* pathogenic variant P525L. However, the mechanisms linking genetic and environmental factors to NMJ defects remain elusive. By taking advantage of co-cultures of motoneurons and skeletal muscle derived from human induced pluripotent stem cells (iPSCs), we show that the neural RNA binding protein HuD (ELAVL4) may underlie NMJ defects and apoptosis in FUS-ALS. HuD overexpression in motoneurons phenocopies the severe FUS^P525L^ mutation, while HuD knockdown in FUS^P525L^ co-cultures produces phenotypic rescue. We validated these findings *in vivo* in a *Drosophila* FUS-ALS model. Neuronal-restricted overexpression of the HuD-related gene, *elav*, produces *per se* a motor phenotype, while neuronal-restricted *elav* knockdown significantly rescues motor dysfunction caused by FUS. Finally, we show that HuD levels increase upon oxidative stress in human motoneurons and in sporadic ALS patients with an oxidative stress signature. On these bases, we propose HuD as an important player downstream of FUS mutation in familial ALS, with potential implications for sporadic ALS related to oxidative stress.

## INTRODUCTION

The neurodegenerative disease amyotrophic lateral sclerosis (ALS) is caused by progressive loss of skeletal muscle function due to motoneuron (MN) death (Brown & Al-Chalabi, 2017). About 10% of the cases are classified as inherited (familial ALS), while the vast majority are considered sporadic. Pathogenic variants in several genes have been linked to both familial and sporadic ALS cases. Most of these variants map in the *C9ORF72* and *SOD1* genes, but several ubiquitously expressed RNA-binding proteins (RBPs) have been linked to ALS. In the nervous system, ubiquitous and neural-specific RBPs cooperate in gene expression regulation and cross-regulation and feedback loops are crucial in establishing proper levels and activity of each RBP (Dassi, 2017). An extensive crosstalk also exists between ubiquitous ALS-linked RBPs, such as FUS, and other neural RBPs expressed in the MN (Blokhuis et al., 2016; De Santis et al., 2019). Thus, in the MN, ALS-associated genetic variants in one node of the network might impact other RBPs and produce broader, unexpected, effects that could be highly cell-type specific. In this context, we have previously shown that mRNA and protein levels of HuD, encoded by the *ELAVL4* gene, are upregulated in MNs derived from *FUS* mutant human induced pluripotent stem cells (iPSCs) and in a mouse model (Fus-Δ14 knock-in) (Garone et al., 2021). Increased HuD levels in FUS^WT^ MNs produce changes in the transcriptome that are substantially similar to those exerted by the severe *FUS* P525L variant (Garone et al., 2023). A direct consequence of HuD upregulation in FUS mutant iPSC-derived MNs and mouse spinal cord is an increase in the levels of its targets, including Neuritin1 (NRN1; also known as cpg15) and GAP43 (Garone et al., 2021). HuD plays key roles during nervous system development and recent evidence points to its possible involvement in neurodegenerative processes as well (Bronicki and Jasmin, 2013; Dell’Orco et al., 2021; Silvestri et al., 2022).

In this work we aimed to assess if HuD dysregulation could underlie neuromuscular junctions (NMJs) defects in the context of ALS. Several genes that are commonly altered in FUS^P525L^ MNs and HuD overexpressing MNs are involved in axon projection, cell junction and synapse formation (Garone et al., 2020; Garone et al., 2023). ALS is considered a distal axonopathy as early impairment of NMJs has been reported in both animal models and patients. These include SOD1^G93A^ mice, in which endplate denervation, followed by ventral root axons degeneration, occurs way before the death of MNs in the spinal cord (Fischer et al., 2004). Notably, such ‘‘dying back’’ pattern was also evident in a sporadic ALS patient who unexpectedly died few weeks after the diagnosis (Fischer et al., 2004), and recently proposed for familial ALS linked to RBP genes, including *FUS*. Increased apoptosis and lower MNs loss due to toxic gain of function of the mutant protein was observed at birth in a knock-in *Fus* mouse model (*Fus^ΔNLS/ΔNLS^*; Scekic-Zahirovic et al., 2016). Notably, MN loss in *Fus^ΔNLS/ΔNLS^*mice occurred in the absence of FUS protein aggregation and stress granules formation and was rescued by MN-specific reversion of FUS mislocalization (Scekic-Zahirovic et al., 2016). These findings point to cell-autonomous mechanisms leading to MN death which are due to the gain of toxic cytoplasmatic functions of the FUS mutant protein and are independent from its aggregation. A direct causal link between *FUS* variants and NMJ defects was demonstrated by Picchiarelli et al., who showed a reduction in the number and area of the endplates in newborn *Fus^ΔNLS/ΔNLS^* mice and progressive denervation and reduction of the endplate size in adult heterozygous mutants (*Fus^ΔNLS/+^*). (Picchiarelli et al., 2019). This phenotype, which was observed in human iPSC-derived co-cultures as well, was due to cell-autonomous toxicity of mutant FUS in both skeletal muscle cells (SkMCs) and MNs. In the muscle, innervation induces FUS enrichment in subsynaptic nuclei, where it promotes expression of *Chrn* genes. However, this cannot be considered the sole underlying mechanism for the observed NMJ defects, as *FUS* mutation in MNs exerted a similar phenotype upon co-culture with wild-type iPSC-derived SkMCs in the absence of any difference in AChR genes expression in the muscle (Picchiarelli et al., 2019). Decreased number of NMJs per myotube and impaired NMJ maturation were also observed in co-cultures of mutant FUS iPSC-derived MNs and human myotubes derived from mesoangioblasts (MAB) from a healthy individual (Stoklund Dittlau et al., 2021).

Here we found that HuD gain-of-function can account for most of the cell-autonomous effects of a severe *FUS* variant in the MN. These include defective maturation of the NMJ and apoptosis, possibly by a “dying back” pattern, in co-cultures of iPSC-derived MNs and SkMCs. These findings were further supported *in vivo* using a Drosophila model, in which reduction of the *HuD*-related gene *elav* is sufficient to rescue the motor phenotype caused by FUS overexpression. Strikingly, in both iPSC-derived co-cultures and Drosophila, HuD (or ELAV) overexpression produces relevant ALS phenotypes *per se*. Thus, we wondered if HuD-dependent mechanisms might underlie sporadic ALS independently of *FUS* mutation. Oxidative stress is thought to play a major role as an environmental factor involved ALS onset and progression (Brown & Al-Chalabi, 2017). Notably, increased HuD levels were recently reported in neuroblastoma cells upon oxidative stress and in the motor cortex from few sporadic ALS (sALS) patients (Dell’Orco et al., 2021). Here, by reanalyzing the data from a large transcriptome study (Tam et al., 2019), we showed that HuD expression is specifically increased only in sALS cases with an oxidative stress signature (accounting for about 60% of the cohort), while it is decreased or unchanged in the other groups. We propose that HuD-dependent pathological mechanism directly contribute to familial FUS-ALS and in a distinct group of sALS cases associated with oxidative stress.

## RESULTS

### Establishment of iPSC-derived neuromuscular co-cultures

We have previously developed methods to obtain iPSC-derived functional MNs and SkMCs (Lenzi et al., 2016; Garone et al., 2019). The protocol for SkMCs generation was improved to reduce the subpopulation of differentiation resistant cells in more extended cultures (Supplementary Fig. S1A-D). Spinal MN progenitors, generated in 5 days as previously described (Garone et al., 2019), were re-plated onto SkMCs obtained in 13 days with the optimized protocol (Figure 1A,B). Recombinant agrin and laminin proteins were added to the culture medium to promote the expression of post-synaptic AChR subunit genes in muscle cells through the activation of the agrin–LRP4–MuSK–ERM axis, and pre-synaptic differentiation and neurotransmitter release, respectively (Ichikawa et al., 2005; Stoklund Dittlau et al., 2021). After this step, hereafter considered day 0 of the co-culture, we maintained MNs and SkMCs for further 50 days. Clustering of acetylcholine receptors (AChRs), detected by α-bungarotoxin (αBTX) staining, suggested the establishment of motor endplates as early as day 14 (Figure 1C). The co-localization of the presynaptic marker SYNAPTOPHYSIN (SYP) and αBTX signals further indicated the establishment of neuromuscular junctions (NMJs) (Figure 1D). At day 50, SkMCs show few spontaneous contractions, which were significantly increased by addition of glutamate, to stimulate MNs, and subsequently reduced to zero by addition of tubocurarine, a non-depolarizing competitive antagonist of the nicotinic AChR at the NMJ (Figure 1E). Notably, SkMCs cultured in the same conditions, but in absence of MNs, never showed contractions.

**Figure 1.**
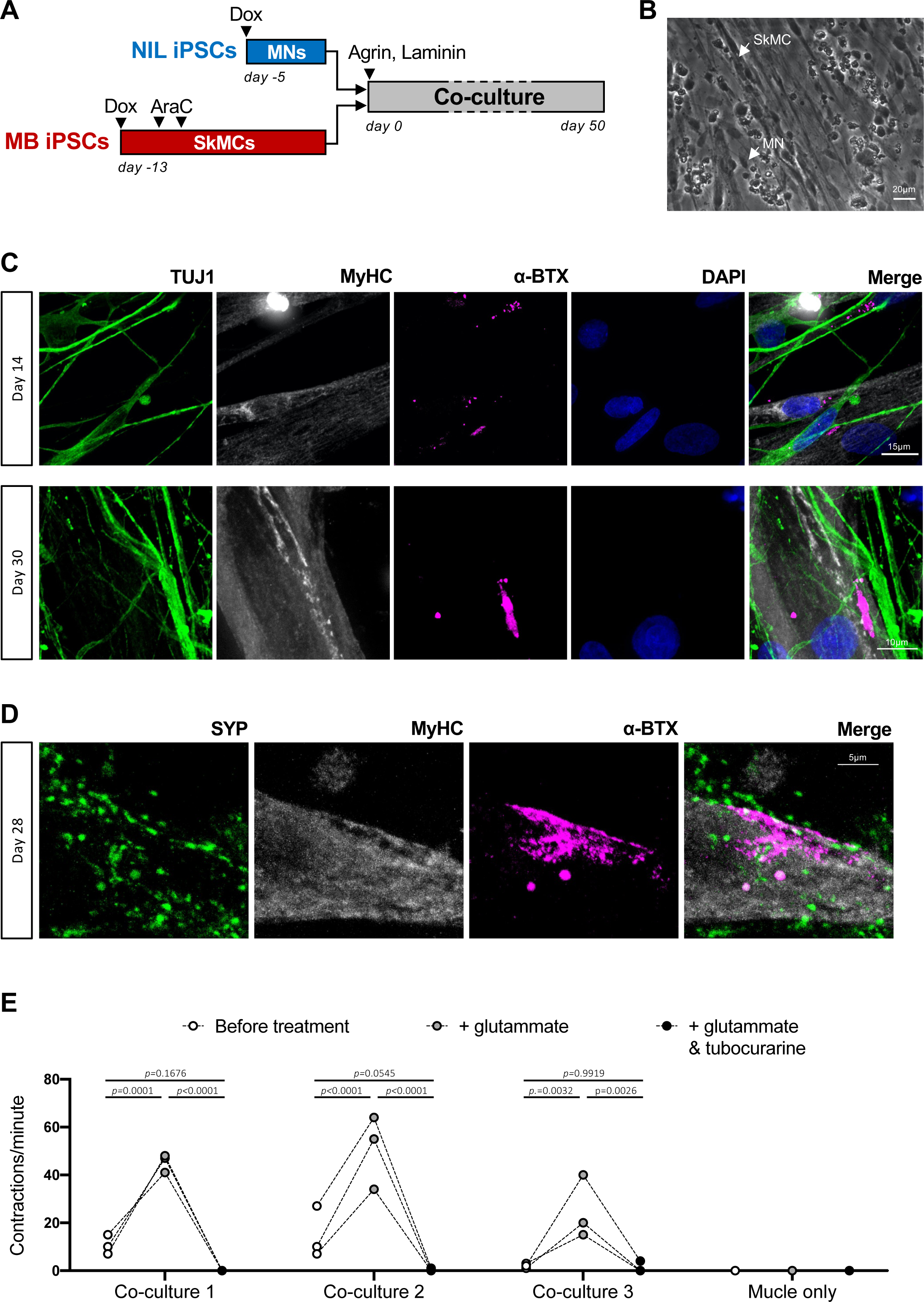
Establishment of iPSCs derived MNs and SkMCs co-cultures and functional analysis of NMJs. (A) Schematic representation of the differentiation protocol to generate co-cultures of spinal MNs and SkMCs. A more detailed SkMC differentiation scheme is shown in Supplementary Figure S1. (B) Phase-contrast image of MNs and SkMCs co-culture derived from FUS^WT^ iPSCs at day 14. Examples of MN cell body and SkMC are indicated by arrows. Scale bar: 20 μm. (C,D) Immunostaining analysis of co-cultures at the indicated time points. α-BTX (magenta) was used as a AChRs clustering marker, TUJ1 (TUBB3, green in panel C) and MyHC (grey) are, respectively, neuronal and muscular markers. SYP (green in panel D) is a presynaptic marker. DAPI (blue) was used for nuclear staining. Scale bar: 15 μm, 10 μm or 5 μm as indicated. (E) The graph shows the number of contractions per minute in day 50 co-cultures before treatment, 5’ upon glutamate addition, and 5’ upon further addition of tubocurarine. Contractions were counted in 3 randomly selected fields (each represented by dots connected by a line) of 3 independent co-cultures. One representative monoculture of SkMCs only, showing no contractions in all conditions, is also displayed. Two-way repeated measures ANOVA (performed only among co-cultures): for treatment comparison *p*<0.0001, for replicate comparison *p*=0.0837; post hoc Tukey test *p* values for multiple comparisons among treatments within each replicate are indicated in the graph.

Collectively, our data suggests a successful development of a functional neuromuscular model system for mechanistic studies.

### NMJ defects due to FUS mutation or HuD overexpression in MNs

Impaired NMJ formation upon FUS mutation in SkMCs, MNs or both had been previously reported (Picchiarelli et al., 2019). We aimed to assess whether the detrimental effect of mutant FUS expression in the MN could be due, at least in part, to increased HuD levels. To this aim, we established co-cultures using MNs from iPSCs with 3 different genetic backgrounds: the isogenic FUS^WT^ and FUS^P525L^ (Lenzi et al., 2015), and FUS^WT^ overexpressing HuD under the Synapsin1 promoter (Syn1::HuD; hereafter indicated as “FUS^WT^+HuD”; Garone et al., 2021) (Figure 2A). Notably, the Synapsin1 promoter confers overexpression of HuD in the range observed in FUS^P525L^ MNs (Garone et al., 2021). In all subsequent experiments, SkMCs have always been derived from FUS^WT^ iPSCs.

**Figure 2.**
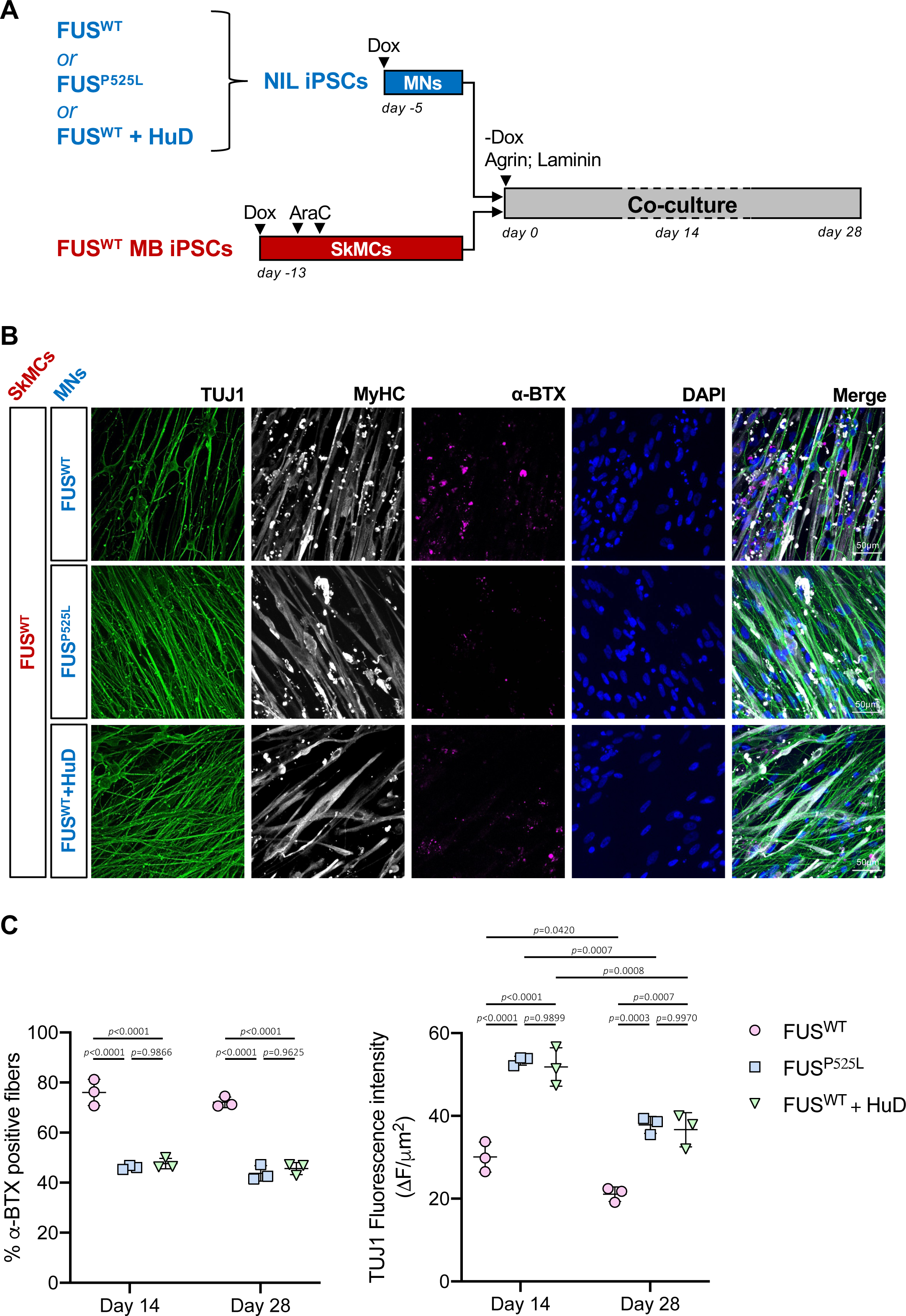
NMJ analysis in co-cultures. (A) Schematic representation of co-cultures used for analyses shown in Figures 2-5. (B) Representative images of immunofluorescence staining of day 14 co-cultures using the indicated primary antibodies and α-BTX and DAPI. Scale bar for all panels: 50 μm. (C) The graphs report quantitative analysis of the percentage of α-BTX positive fibers (left) and TUJ1 fluorescence intensity (right), at day 14 and 28 of co-culture. Each dot represents a replicate, consisting of an individual batch of differentiated iPSCs. For each replicate, 4-6 randomly selected fields were used for α-BTX positive fibers analysis, and 6 randomly selected fields for TUJ1 analysis. Error bars indicate standard deviation calculated on the average value of the replicates. Ordinary two-way ANOVA: for α-BTX positive fibers, *p*=0.0715 between day 14 and 28, *p*<0.0001 among genotypes; for TUJ1 fluorescence intensity, *p*<0.0001 for both time points and genotypes comparisons; relevant post hoc Tukey test *p* values for multiple comparisons among genotypes within each time point are indicated in the graph.

After 14 and 28 days, we observed reduction in muscle fibers showing AChR clusters, detected by αBTX staining, in the co-cultures established with FUS^P525L^ MNs or upon HuD overexpression (Figure 2B,C). According to our previous observations (Garone et al., 2021), both FUS mutant and HuD overexpressing MNs showed increased neurite network (Figure 2B,C). Thus, the reduction of skeletal muscle cells forming endplates cannot be explained by a reduction in the neuronal component in FUS mutant, or HuD overexpressing, MN co-cultures. Detailed analysis of the endplate morphology suggested defective maturation when mutant FUS was expressed in the MN, confirming previous observations (Picchiarelli et al., 2019). In particular, we observed a reduced fraction of aligned AChR puncta and endplates categorized as “dense” or “clustered” (Figure 3A,B). Conversely, endplates with diffuse AChR puncta, indicative of an immature formation stage, were significantly increased. Notably, also in this case, HuD upregulation in an otherwise FUS WT genetic background perfectly phenocopied the defects in NMJ maturation produced by the FUS P525L variant.

**Figure 3.**
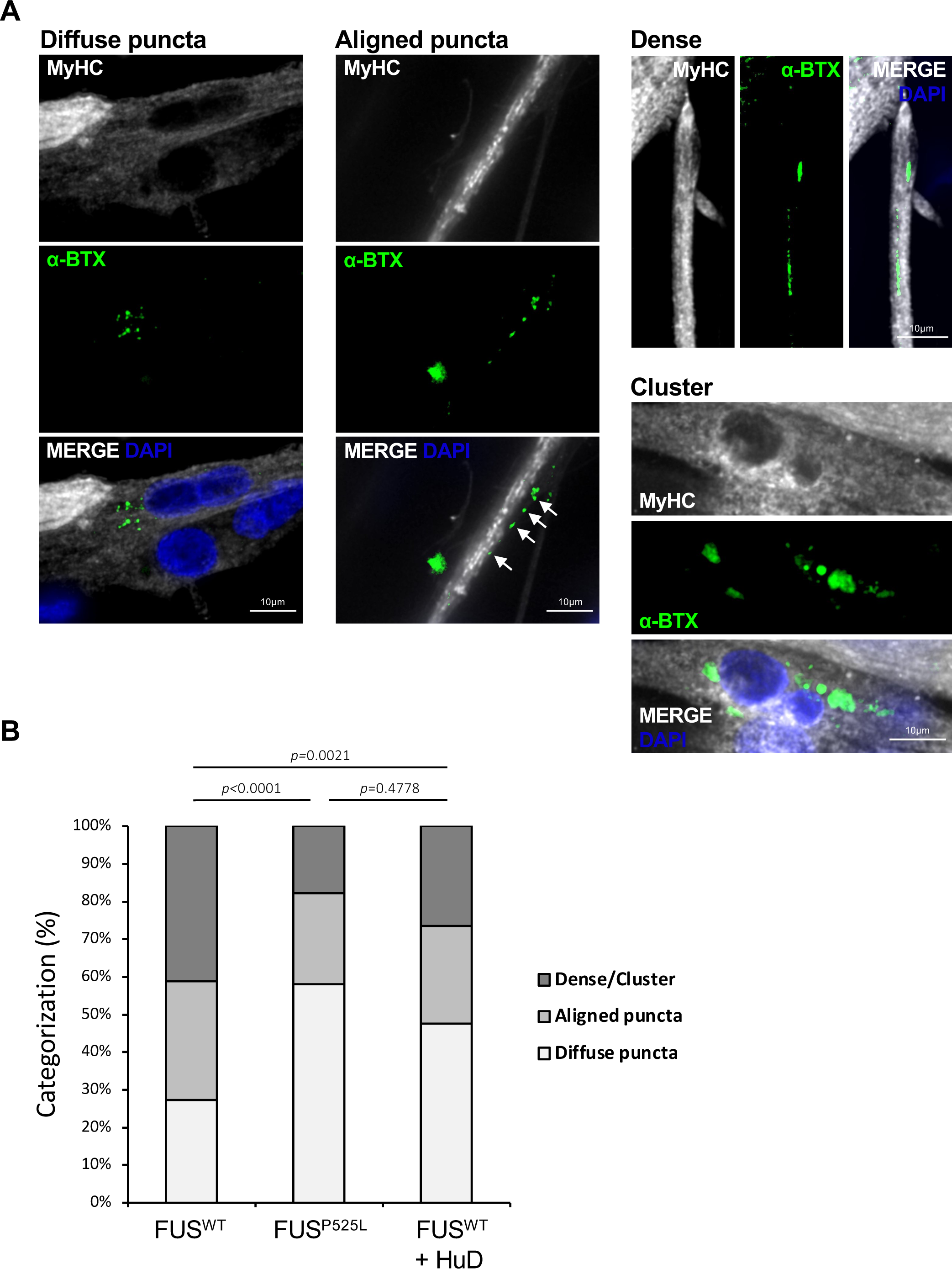
Endplate maturation analysis in co-cultures. (A) Representative images of immunofluorescence staining of day 14 co-cultures using the MyHC antibody and α-BTX and DAPI. Images show representative endplates that were categorized based on morphology into diffuse puncta, aligned puncta (indicated by arrows) and dense/cluster. Scale bar: 10 μm. (B) The graph shows quantification of endplates categories, as a percentage, from co-cultures obtained with FUS^WT^ SkMCs and FUS^WT^, FUS^P525L^ or FUS^WT^+HuD MNs. The results of a blind analysis of 165 (FUS ^WT^), 124 (FUS^P525L^), and 147 (FUS^WT^+HuD) total endplates from 3 batches of differentiated cells are shown. Fisher’s exact test with Bonferroni correction for multiple testing.

### FUS mutation or HuD overexpression in MNs induced cell death by apoptosis upon co-culture with SkMCs

At late time points, beyond day 28, extensive death of both neuronal and muscle cells was observed in co-cultures containing FUS^WT^ SkMCs in presence of FUS^P525L^ or FUS^WT^+HuD MNs (Fig. 4A). Quantification of dead cells by staining with fluorescent ethidium homodimer-1 in a time course analysis revealed, for both genotypes, significant difference compared to the FUS^WT^ control (Fig. 4B; Supplementary Figure S2). Notably, we did not find any significant difference between co-cultures made by FUS^P525L^ and FUS^WT^+HuD MNs. Within each line, a significant increase of dead cells from the initial time point (day 7) was observed for the FUS^WT^ co-cultures only at day 28, while it was evident at earlier time points for FUS^P525L^ (from day 14 onward) and FUS^WT^+HuD (from day 21 onward) (Fig. 4B, p values table). Increased Cleaved Caspase-3 (CC-3) signal at day 14 and 28 suggests that cell death was a consequence of apoptosis (Figure 4C-E; Supplementary Figure S3). From day 14 to day 28, the number of apoptotic cells significantly increased in the co-cultures made by FUS^P525L^ and FUS^WT^+HuD, but not FUS^WT^, MNs (Figure 4E). Importantly, when MNs are cultured in absence of muscle cells, we did not observe differences in CC-3 signal among the 3 genotypes (Figure 4D).

**Figure 4.**
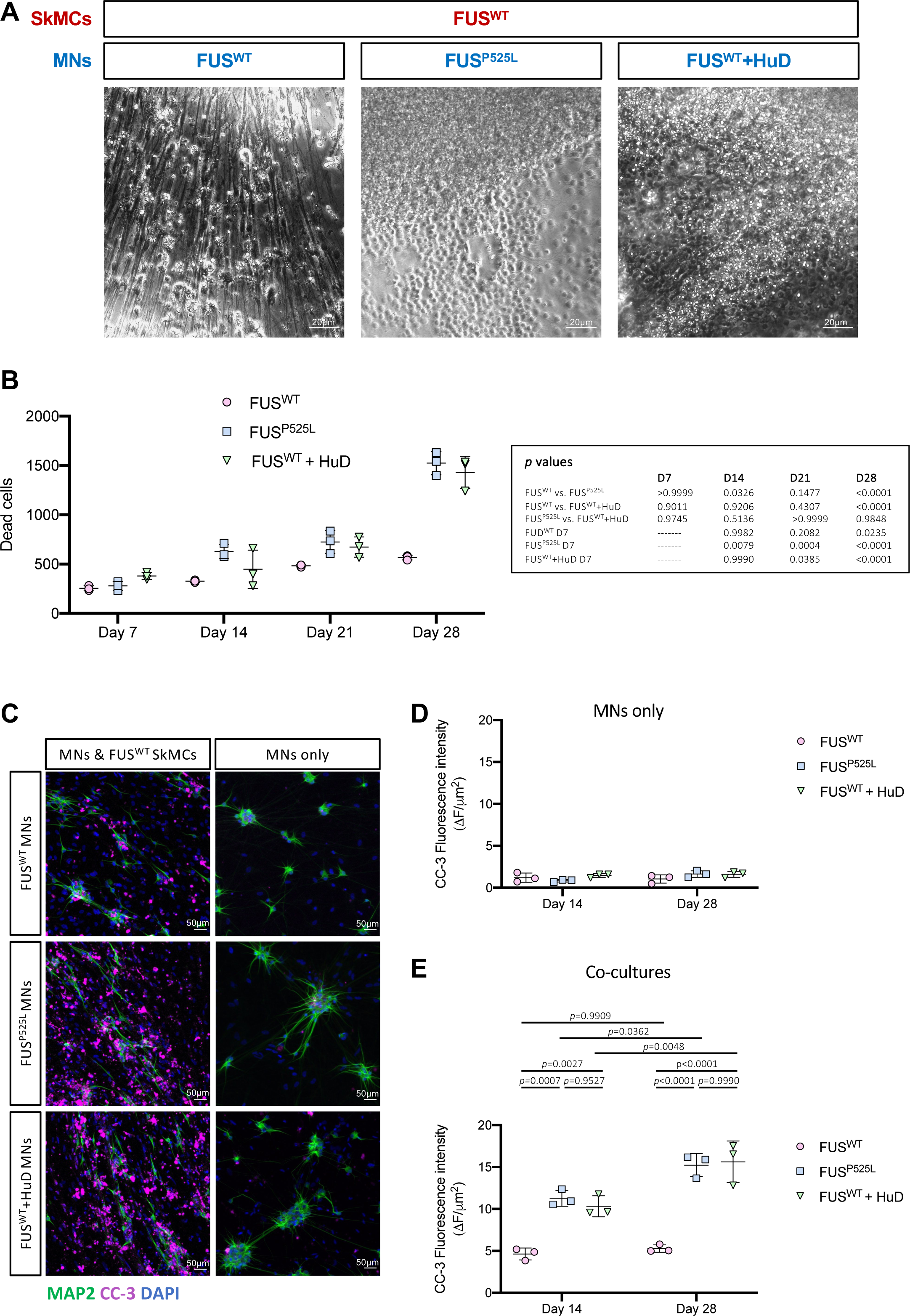
Cell death analysis. (A) Representative brightfield images of co-cultures at day 28. Scale bar: 20μm. (B) Quantitative analysis of dead cells using ethidium homodimer-1 at day 7, 14, 21, 28 of co-cultures obtained with FUS^WT^ SkMCs and FUS^WT^, FUS^P525L^ or FUS^WT^+HuD MNs. The graph shows the average (dots) and standard deviation from 3 batches of differentiated cells. For each replicate, 5 fields have been acquired at each time point. Ordinary two-way ANOVA: *p*<0.0001 for both time points and genotypes comparisons and their interaction; post hoc Tukey test *p* values for multiple comparisons among genotypes within each time point, and among day 7 and later time points within each genotype, are indicated in the table. See Supplementary Figure S2 for representative images of the cells. (C) Representative merged images of immunofluorescence staining of day 14 co-cultures (left panels) or MN monocultures (right panels) using Cleaved Caspase-3 (CC-3) and MAP2 antibodies, and DAPI. Scale bar: 50 μm. Single panels are shown in Supplementary Figure S3. (D,E) The graphs report quantitative analysis of the CC-3 fluorescence intensity at day 14 and 28 of MNs monoculture (D) or co-cultures (E). Each dot represents a replicate, consisting of an individual batch of differentiated iPSCs. For each replicate 6 fields have been acquired and the dot shows the average value. Error bars indicate standard deviation calculated on the average value of the replicates. Ordinary two-way ANOVA. No significant differences resulted from the comparison of MN monocultures (D). For the co-cultures (E), *p*<0.0001 for genotypes comparison, *p*=0.0003 for time points comparison, *p*=0.0324 for their interaction; relevant post hoc Tukey test *p* values for multiple comparisons are indicated in the graph.

To confirm these observations in an independent pair of isogenic FUS lines, considered as a biological replicate, we introduced the P525L variant in another iPSC line (EBiSC catalogue number WTSIi004-A, male, age of donor 35-39) (Supplementary Figure S4A). In these lines, hereafter named WTSI-FUS^WT^ and WTSI-FUS^P525L^, we confirmed upregulation of HuD in FUS mutant MNs (Supplementary Figure S4B). Moreover, when co-cultured with FUS^WT^ SkMCs, we observed reduction in muscle fibers forming endplates and increased apoptosis in presence of WTSI-FUS^P525L^ MNs, compared with the co-cultures obtained with isogenic WTSI-FUS^WT^ MNs (Supplementary Figure S4C,D; Supplementary Figure S5A,B). We then produced a WTSI-FUS^WT^ line containing the Syn1::HuD transgene (hereafter WTSI-FUS^WT^+HuD). Also in this case, HuD overexpression resulted in a significant reduction of muscle fibers forming NMJs and increase in apoptosis (Supplementary Figure S4C,D; Supplementary Figure S5A,B).

Collectively, these results show that, in addition to NMJ formation impairment, HuD overexpression in MNs closely phenocopies the severe FUS P525L ALS variant in terms of increased vulnerability by apoptosis, and that such effect emerges only in presence of muscle cells.

### Knockdown of HuD or its target genes in FUS^P525L^ co-cultures rescues NMJ defects and apoptosis

Since the effects of HuD overexpression in MNs are strikingly similar to those exerted by the FUS^P525L^ variant, we wondered whether reduction of HuD levels could be beneficial for NMJ formation and survival in co-cultures made by FUS^P525L^ MNs. Small interfering RNAs (siRNAs) transfection effectively reduced *HuD* mRNA and protein levels in FUS^P525L^ MNs (Supplementary Figure S6A,B). FUS^WT^ SkMCs and FUS^P525L^ MNs co-cultures were transfected at day 3 and 8 with anti-HuD siRNAs and analyzed at day 14 by αBTX and CC-3 staining. This analysis revealed significant increase in the fraction of muscle fibers showing AChR clusters and significant reduction of apoptosis upon HuD knockdown (Figure 5A-D; Supplementary Figure S6C,D). The same effects were consistently observed upon HuD siRNAs transfection in co-cultures made of WTSI-FUS^P525L^ MNs (Supplementary Figure S7A,B). We next performed a similar experiment by knocking down, individually, two key HuD targets, *NRN1* and *GAP43*, whose levels are increased in FUS^P525L^ MNs due to overly stabilization by HuD (Supplementary Figure S6A; Garone et al., 2021). *NRN1* siRNAs significantly increased the fraction of muscle fibers showing AChR clusters and decreased apoptosis to a similar extent as *HuD* siRNAs (Figure 5A-D; Supplementary Figure S6C,D). *GAP43* knockdown did not significantly change AChR clustering and produced a milder effect on apoptosis.

**Figure 5.**
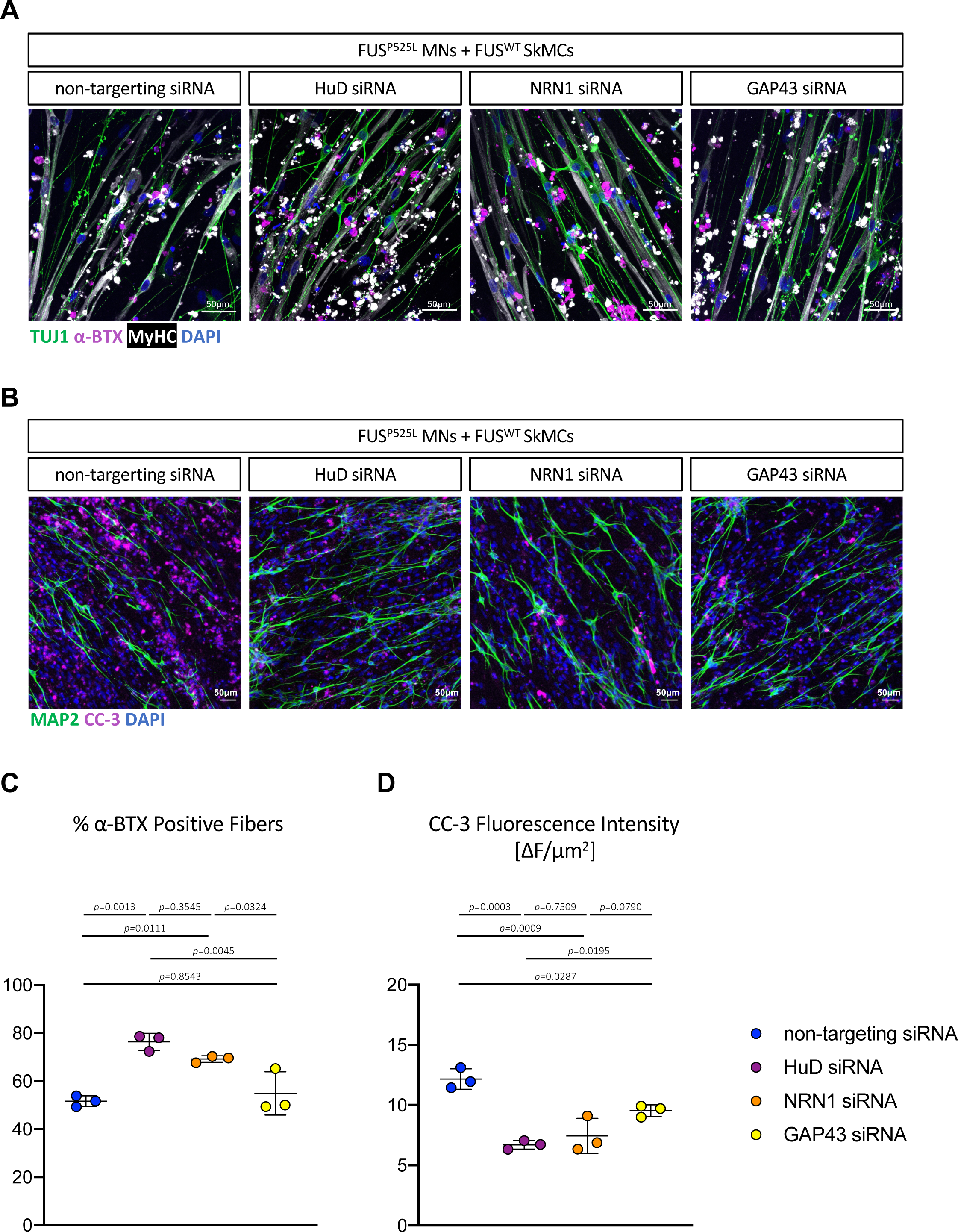
HuD, NRN1 and GAP43 knockdown in co-cultures containing FUS^P525L^ MNs. (A,B) Representative merged images of day 14 co-cultures obtained with FUS^WT^ SkMCs and FUS^P525L^ MNs transfected with the indicated siRNAs and stained with the indicated antibodies. Scale bar: 50μm. Single panels are shown in Supplementary Figure S6. (C,D) The graphs report quantitative analysis of the percentage of α-BTX positive fibers (C) and CC-3 fluorescence intensity (D), at day 14 of co-culture as in (A,B). Each dot represents an individual batch of differentiated iPSCs transfected with the indicated siRNAs. For each replicate, 5-7 fields have been acquired for α-BTX positive fibers analysis and 8-10 fields for CC-3 immunofluorescence. The dot shows the average value. Error bars indicate standard deviation calculated on the average value of the replicates. Ordinary one-way ANOVA, *p*=0.0009 for α-BTX positive fibers, *p*=0.0003 for CC-3 immunofluorescence; post hoc Tukey test *p* values for multiple comparisons are indicated in the graph.

Taken together these results suggest that most, if not all, the cell-autonomous effects of the FUS^P525L^ variant in MNs in terms of NMJ defects and apoptosis could be due to HuD upregulation. In turn, among the tested HuD targets that are also upregulated in FUS mutant MNs (Garone et al., 2021), NRN1 might play a prominent role.

### Knockdown of the Drosophila HuD ortholog *elav* rescues the motor dysfunction caused by FUS overexpression *in vivo*

Drosophila models represent valuable tools for examining genetic modifiers of neurodegenerative diseases, including FUS-ALS (Casci and Pandey, 2015). In *Drosophila melanogaster*, the neuronal *embryonic lethal abnormal vision* (*elav*) gene encodes for an RBP, ELAV, showing high conservation with the mammalian HuD protein in terms of structure, expression and functions (Bronicki and Jasmin, 2013). Upon ELAV overexpression in neurons, we observed a strong reduction in the number of flies able to cross a 4cm threshold in a climbing assay performed at day 6 and 15 (Figure 6A). We took advantage of fly models selectively overexpressing human FUS WT or P525L in neurons (Supplementary Figure 8A), which show reduced climbing ability (Marrone et al., 2019), to assess whether *elav* modulation might modify such motor dysfunction phenotype. In particular, we attempted a rescue experiment by ELAV reduction by neuronal-specific expression of short hairpin RNAs (shRNAs). We did not observe any difference at day 6, an early age at which neither WT nor mutant FUS overexpression produced motor dysfunction (near 100% of flies able to climb 4cm in 10s). Strikingly, *elav* RNAi significantly rescued the motor dysfunction for both WT and P525L FUS at day 15 (Figure 6B). This effect was further confirmed by analyzing a different motor parameter, climbing velocity, which was significantly improved by ELAV reduction in both WT and P525L FUS at day 15 (Figure 6C). We then analyzed the percentage of mature and collateral boutons in third instar larvae, as an indication of the maturity of NMJs (Figure 6D; Supplementary Figure 8B). As shown in Figure 6E, *elav* RNAi reduced the percentage of collateral boutons and increased the percentage of mature boutons in FUS WT expressing larvae. However, we did not observe a similar effect in mutant FUS expressing larvae. An opposite trend, reduced mature boutons and increased collateral boutons, was observed in larvae overexpressing ELAV, however the differences did not reach statistical significance (Supplementary Figure 8C).

**Figure 6.**
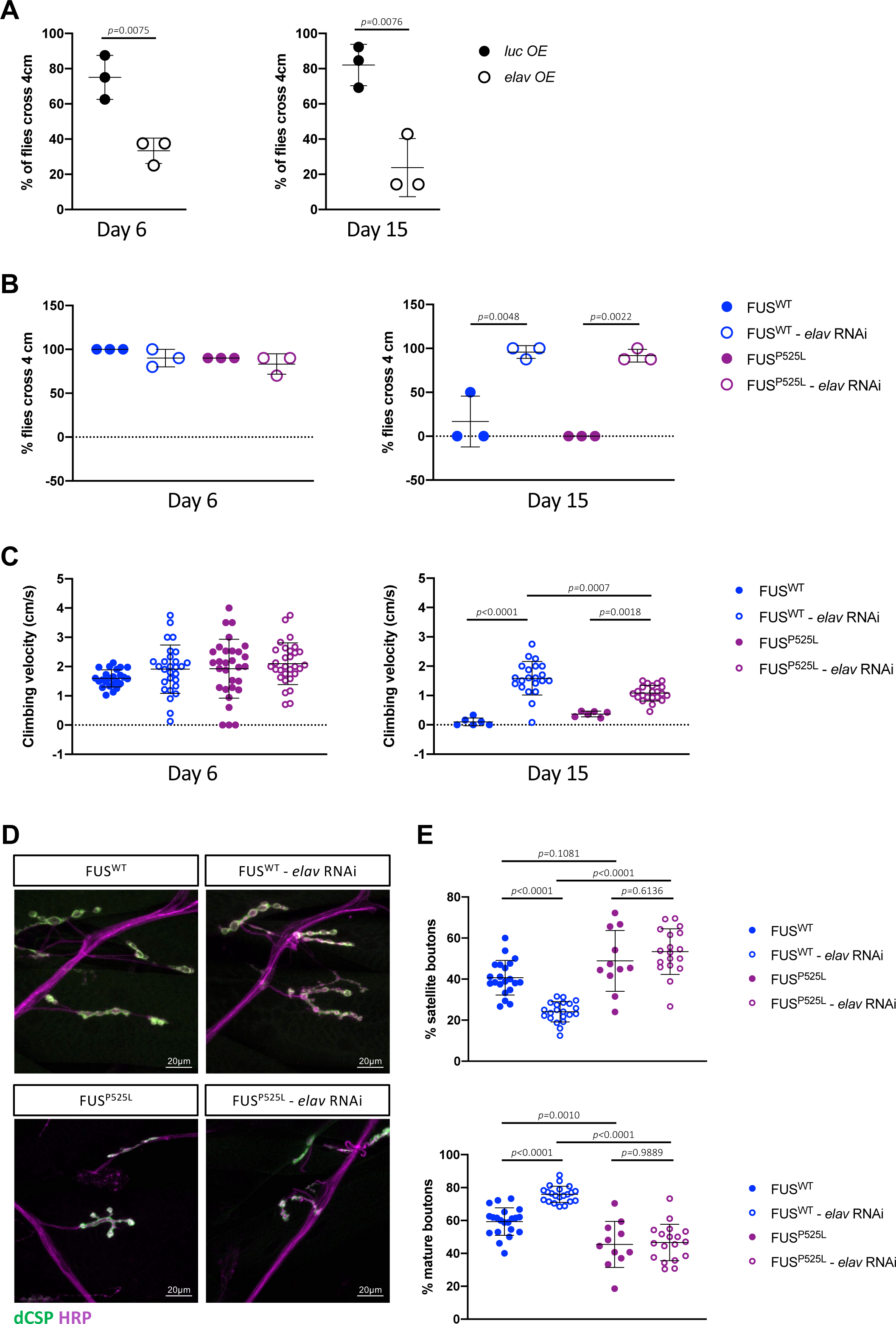
Expression of *Drosophila* HuD ortholog *elav* modifies FUS-mediated motor dysfunction. (A) The graphs represent the percentage of flies that climb 4cm in 10s at day 6 and day 15 in overexpressing *elav* (*elav* OE) or Luciferase (LUC OE) as a control, respectively. N=3 experimental replicates (trials) with 6-10 flies each. Student’s t-test, unpaired, two tails, *p* values are indicated in the graphs. (B) The graphs show percent flies that climb 4cm in 10s at day 6 and day 15 in wild-type FUS (FUS-WT) or mutant FUS (P525L) with or without *elav* RNAi. N=3 experimental replicates (trials) with 6-10 flies each. Ordinary one-way ANOVA, relevant post hoc Tukey test *p* values for multiple comparisons are indicated in the graphs. (C) The graphs show the climbing velocity (cm/s) of flies as in (B). Each data point represents one fly. Ordinary one-way ANOVA, relevant post hoc Tukey test *p* values for multiple comparisons are indicated in the graphs. (D) Immunohistochemistry analysis of the neuronal marker horseradish peroxidase (HRP) and the synaptic vesicle marker cysteine string protein (dCSP) in third instar larvae expressing FUS-WT or FUS-P525L, with or without *elav* RNAi. (E) The graphs show quantitative analysis of the percentage of satellite and mature boutons in larvae as in (D). N=6-10 larvae per genotype. Ordinary one-way ANOVA, relevant post hoc Tukey test *p* values for multiple comparisons are indicated in the graphs. In all graphs, bars indicate the mean and standard deviation,

Collectively, in line with the results from the *in vitro* human model, these *in vivo* experiments in *Drosophila* show that overexpression of the HuD-related protein ELAV produces *per se* motor dysfunction, whereas ELAV dampening is sufficient to rescue the motor phenotype produced by FUS completely.

### HuD might be involved in sporadic ALS with an oxidative stress signature

Our previous work showed increased HuD levels in FUS mutant human iPSC-derived MNs and in the spinal cord of a FUS mutant mouse model (Garone et al., 2021). We and others have also previously detected HuD in pathological aggregates of FUS, TDP-43 and sALS patients (Blokhuis et al., 2016; De Santis et al., 2019). More recently, the Cereda lab reported HuD overexpression in sALS patients’ motor cortex and proposed that this could be due to induction of *HuD* expression upon oxidative stress, as *HuD* mRNA levels were increased in neuroblastoma cells treated with H_2_O_2_ (Dell’Orco et al., 2021). These findings prompted us to assess whether the same effect could be observed in human MNs. MNs from FUS^WT^ iPSCs were treated with sodium arsenite, an inducer of oxidative stress, for 60 minutes before mRNA analysis by qRT-PCR. We found a small but highly reproducible increase in *HuD* transcript levels upon oxidative stress (Figure 7A). We observed a similar effect in FUS^WT^ iPSC-derived co-cultures (Figure 7B). Upregulation of HuD was likely due to new transcription, as pre-treatment with α-amanitin, a potent and selective inhibitor of RNA polymerase II, abolished the effect of oxidative stress on this transcript (Figure 7C). All these results were confirmed in WTSI-FUS^WT^ MNs and co-cultures (Figure 7D-E). Together with the previous findings (Dell’Orco et al., 2021), these results suggest that oxidative stress might trigger *HuD* expression increase in human neurons, including MNs, with possible implications for sALS. To investigate this possibility, we took advantage of a large transcriptome profiling dataset of sALS patients (Tam et al., 2019). In this dataset, ALS patients’ cortex samples could be grouped into 3 distinct clusters based on expression profile. The largest group included patients with hallmarks of oxidative and proteotoxic stress (ALS_Ox). The other two clusters showed signatures of glial activation (ALS_Glia) and high retrotransposon expression (ALS_TE) (Tam et al., 2019). Notably, we found that *HuD* expression was significantly increased only in the ALS_Ox cluster (Figure 7G). Conversely, its levels were significantly lower in the ALS_Glia group and no significant change was found in ALS_TE group. This pattern was also evident for the HuD targets, *NRN1* (albeit its increase in ALS_Ox did not reach statistical significance) and *GAP43* (Figure 7G). Moreover, the ELAV family member *ELAVL2*, encoding for an RBP (HuB) closely related to HuD, also showed significant increased expression in ALS_Ox, decreased expression in ALS_Glia and no change in ALS_TE (Supplementary Figure S9A). Finally, *SOD1* mRNA, which is a target stabilized by HuD upon oxidative stress (Dell’Orco et al., 2021), was also significantly upregulated in the ALS_Ox group (Supplementary Figure S9A). In order to gain insights into a possible correlation between the levels of *HuD* (and its targets) and *FUS* in sporadic ALS patients, we analyzed their expression at individual patient level for all groups, including healthy individuals (CTR) and patients with other neurodegenerative diseases (OND). A matrix reporting the Spearman’s correlation coefficients calculated between the expression levels of *FUS* and those of *HuD*, *NRN1* and *GAP43* is shown in Figure S9B. This analysis suggests that there is no correlation between *FUS* and *HuD* expression levels in the OND and ALS_Ox groups, low positive correlation in CTR and ALS_TE groups, and low negative correlation in the ALS_Glia group. For what concerns *NRN1* and *GAP43*, we observed positive correlation with *FUS* in all groups except for ALS_Glia. Finally, when CTR and ALS groups were merged, we observed no correlation for *FUS* and *HuD*, and low positive correlation between *FUS* and *NRN1*, and between *FUS* and *GAP43*. Collectively, the correlation analysis suggests that altered FUS expression is unlikely the underlying mechanism for *HuD* upregulation in patients with an oxidative stress signature. These findings suggest that, in the absence of pathogenic FUS ALS variants, oxidative stress can trigger HuD upregulation in the context of human MNs *in vitro*. Accordingly, HuD levels are specifically increased in sALS patients with a signature related to oxidative stress.

**Figure 7.**
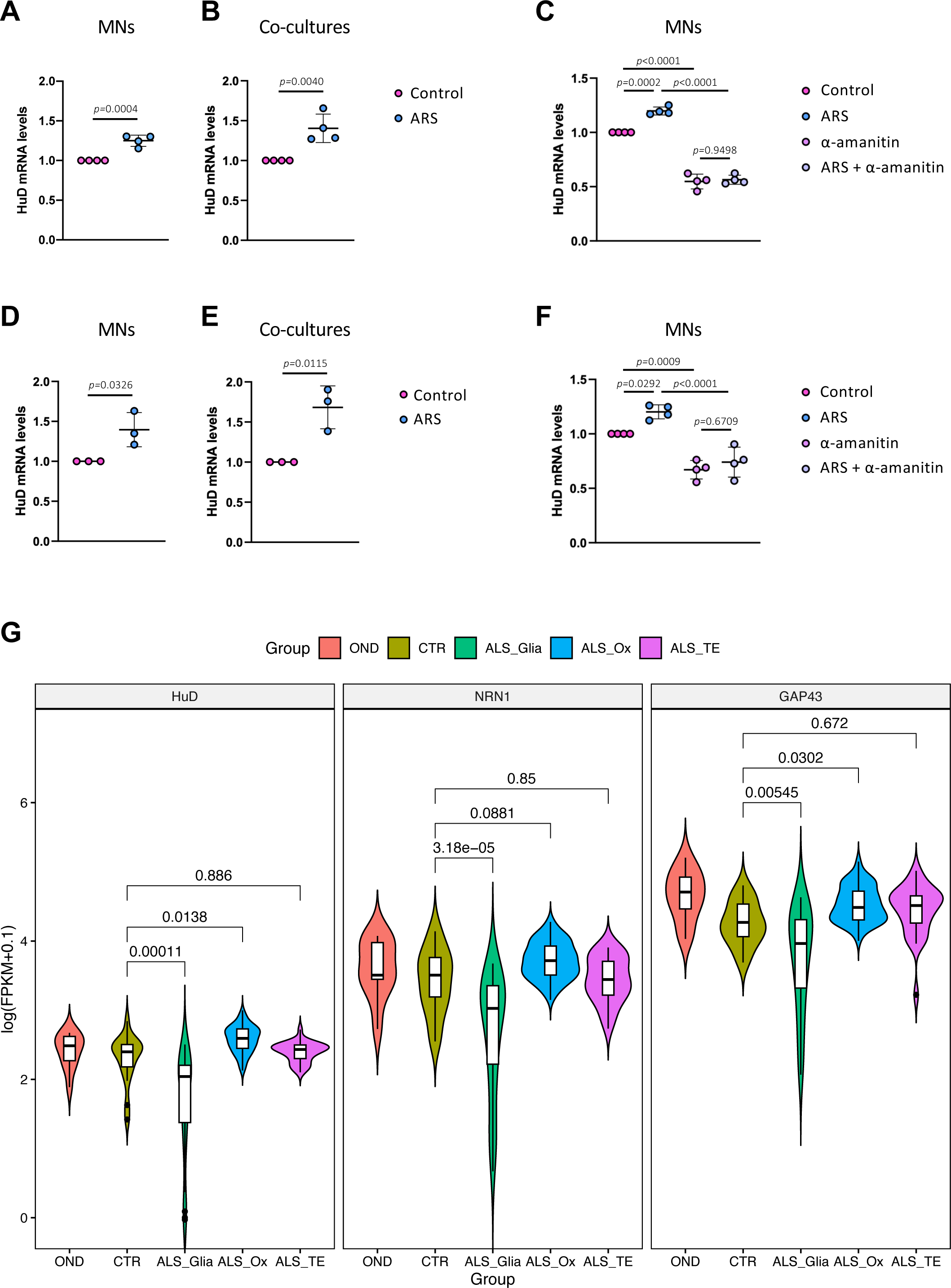
*HuD* expression upon oxidative stress *in vitro* and in sporadic ALS patients. (A,B) The graphs show quantitative analysis of *HuD* mRNA levels in control conditions or upon acute stress treatment with 0.5 mM sodium arsenite (ARS) for 1h in FUS^WT^ iPSCs derived spinal MNs monocultures at day 12 (A) or FUS^WT^ iPSCs derived co-cultures at day 7 (B). (C) Cells as in panel (A) were pre-treated or not with α-amanitin before oxidative stress induction with ARS. (D-F) The experiments shown in panels (A-C) were repeated with the WTSI-FUS^WT^ iPSC line. In panels (A-F) ATP5O expression was used for normalization and the graphs show the average, standard deviation and *p* values from 3 or 4 replicates, each consisting of a batch of differentiated iPSCs. Panels A,B,D,E: Student’s t-test, unpaired, two tails, *p* values are indicated in the graphs. Panels C,F: ordinary one-way ANOVA, *p*<0.0001; relevant post hoc Tukey test *p* values for multiple comparisons are indicated in the graphs. In all graphs, bars indicate the mean and standard deviation, (G) Violin plots showing the expression levels, reported as log-transformed FPKM values, of *HuD*, *NRN1* and *GAP43* in post-mortem sporadic ALS patients’ cortex samples from (Tam et al., 2019). OND: other neurodegenerative diseases; CTR: healthy individuals; ALS_Glia: sporadic ALS patients with a signature of glial activation; ALS_Ox: sporadic ALS patients with a signature of oxidative and proteotoxic stress; ALS_TE: sporadic ALS patients with a signature of high retrotransposon expression. The adjusted *p* values obtained from differential expression analyses are shown.

## DISCUSSION

Human *in vitro* models of the motor unit have shown that MN-restricted expression of ALS-linked FUS variants produces NMJ phenotypes (Picchiarelli et al., 2019; Stoklund Dittlau et al., 2021). Here we found that such NMJ defects could be reverted by dampening HuD levels. Moreover, knockdown of HuD decreased mutant FUS-induced apoptosis *in vitro* and, strikingly, knockdown of the related *elav* Drosophila gene completely rescued motor phenotypes induced by FUS *in vivo*. We have recently shown that in MNs FUS mutation or HuD overexpression exert similar effects on the expression of disease-linked genes (Garone et al., 2023). Altogether, these findings point to a crucial role for HuD in the cell-autonomous mechanisms that underlie the death of MNs via a “dying back” pattern in FUS and, possibly, a subset of sporadic ALS cases with hallmarks of oxidative and proteotoxic stress. To this regard, in both the *in vitro* and the *in vivo* model, we found that HuD (or ELAV) overexpression *per se* phenocopies the severe FUS mutation.

The seminal discovery of early defects at the NMJ in 47-days old SOD1^G93A^ mice, in which loss of MNs occurs only after 90 days of age, supported the “dying back” hypothesis which was further reinforced by evidence from a sALS patient and other non-SOD1 ALS animal models (Fischer et al., 2004; Alhindi et al., 2021). These include the *Fus^ΔNLS/+^* mice that show reduced endplate area and NMJ denervation as early as 1 month of age, while motor deficits begin around 10 months and MN loss was evident only at 22 months (Scekic-Zahirovic et al., 2017; Picchiarelli et al., 2019). Neuron-specific expression of mutant human FUS proteins driven by the *MAPT* regulatory sequences also resulted in early structural and functional defects at the NMJ that preceded MN loss (Sharma et al., 2016). These phenotypes were particularly severe in the MAPT-hFUS^P525L^ mouse, which showed significant denervation at day 20 and MN loss at day 30. This evidence suggests that MN-restricted expression of mutant FUS is sufficient *in vivo* to trigger early retraction of motor axons from the NMJ. Since HuD upregulation was observed in mice carrying a severe FUS mutation in the NLS (Fus-Δ14 knock-in) (Garone et al., 2021), we can speculate that the HuD-dependent circuitry described here could contribute to such NMJ defects due to pre-synaptic mechanisms. For what concerns the Drosophila model used here, the motor phenotype rescue observed upon *elav* RNAi in neurons of FUS^WT^ overexpressing flies was mirrored by a shift towards more mature NMJs. However, we did not observe a significant change of NMJ types in flies overexpressing mutant FUS, suggesting that other mechanisms might contribute to the beneficial effect of ELAV reduction *in vivo*.

The evidence of HuD upregulation in the motor cortex of sporadic ALS patients, possibly due to exposure to environmental factors causing oxidative stress (Dell’Orco et al., 2021), suggests that this RBPs might play a broader role in ALS. To this regard, we and others have previously found HuD in cytoplasmic inclusions of FUS, TDP-43 and sporadic ALS patients (Blokhius et al., 2016; De Santis et al., 2019). In this work we further explored the possibility that HuD might be altered in sporadic ALS. We found increased levels of HuD and some of its targets in a subset of sporadic ALS patients characterized by a signature related to oxidative stress, who account for the largest subgroup (about 60% of the cohort) (Tam et al., 2019). Intriguingly, a significant decrease was instead observed in the cluster with a signature related to glial activation and inflammation (about 20% of the cohort). Thus, taken together, altered HuD levels in either direction would be associated with most sporadic ALS cases. So far, to the best of our knowledge, no known genetic variants in the *HuD* gene have been conclusively linked to ALS. However, from a survey of the Project MinE (http://databrowser.projectmine.com) we noticed variants of uncertain significance (VUS) found in ALS patients but not in controls (Supplementary Figure 10A,B). Future work will be needed to clarify if such VUS have any effect on HuD protein levels and/or activity. The molecular mechanisms underlying *HuD* upregulation upon oxidative stress are unknown. The fact that *HuD* mRNA levels increase upon either arsenite (present work) or H_2_O_2_ treatment (Dell’Orco et al., 2021) point to eIF2α-dependent and -independent pathways (Emara et al., 2012). While regulation of *HuD* by FUS occurs at the post-transcriptional level (De Santis et al., 2017; De Santis et al., 2019; Garone et al., 2021), here we show that *HuD* mRNA increase upon oxidative stress is likely due to transcriptional regulation. A mechanistic link with FUS is therefore unlikely in this latter case. Moreover, we did not find any correlation between *FUS* and *HuD* expression at individual patients’ level in the oxidative stress cluster. Thus, altogether, our previous findings (De Santis et al., 2017; De Santis et al., 2019; Garone et al., 2021) and present work point to FUS-dependent (in FUS-ALS) and FUS-independent (in sALS) mechanisms leading to *HuD* dysregulation in ALS.

Among known HuD targets, in this work we focused on NRN1 and GAP43. Notably, in postmortem ALS patients both GAP43 and NRN1 have been found upregulated in regions of the nervous system with implications in ALS and frontotemporal lobar degeneration (FTLD), including the anterior horn of the spinal cord and the frontal cortex area 8 (Parhad et al., 1992; Ikemoto et al. 1999; Andrés-Benito et al., 2017). This is in line with the known role of HuD in stabilizing (and localizing) *GAP43* and *NRN1* mRNAs through AU-rich elements (ARE) (Wang et al., 2011; Yoo et al., 2013). We found that knockdown of NRN1 or GAP43 produced a partial rescue of NMJ and/or apoptosis phenotypes in co-cultures containing FUS mutant MNs. This effect might be due to their known roles in developing neurons. To this regard, aberrant axon guidance is among the proposed mechanisms underlying the “dying back” phenomenon (Pasterkamp and Giger, 2009; Dadon-Nachum et al., 2011). For instance, it has been shown that increased expression of the axon guidance protein Semaphorin 3A (Sema3A) by terminal Schwann cells (tSCs) might result in axonal denervation at the NMJ due to repulsion of the motor axon in SOD1^G93A^ mice (de Winter, 2006). Notably, GAP43 can modulate axon guidance signals triggered by Sema3A, and exposure to Sema3A induced neuronal death by apoptosis in a GAP43-dependent manner (Gagliardini et al., 2000). Accordingly, loss of GAP43 conferred resistance to apoptotic stimuli in cultured mouse neurons. On the other hand, overexpression of GAP43 in adult mice resulted in prolonged sprouting and death of MNs (Aigner et a., 1995; Harding et al., 1999). In line with these studies in animal models, we show here that knockdown of GAP43 by RNAi partially reverts apoptosis in iPSC-derived co-cultures made by mutant FUS MNs. In this case, however, we speculate that increased HuD levels would mediate GAP43 gain of function in a cell-autonomous manner, rather than by Sema3A signaling, as our in vitro model does not contain tSCs.

We found that NRN1 knockdown in FUS mutant MNs increased AChR clusters. Previous literature on a possible role for NRN1 in NMJ formation is scarce. A relevant study in *Xenopus laevis* tadpoles showed that overexpression of the ortholog gene *cpg15* promoted spinal MN axon growth by increasing branch addition and maintenance and decreasing branch retraction (Javaherian and Cline, 2005). This dysregulation could eventually impact some crucial processes such as extension, refinement, and pruning. Indeed, in such developmental context, new branches formed upon *NRN1*/*cpg15* overexpression were apposed to AChR clusters. However, the authors did not provide information on further maturation of the new endplates, and whether increased branching during development would be followed by productive formation of mature NMJs remains unclear. It should be noted that the adult human spinal cord is among the nervous system districts with the lowest expression of *NRN1* (Garone et al., 2021). We might speculate that aberrantly increased levels of this growth promoting factor might be detrimental, rather than beneficial, in the adult motor system, generating excess of axonal extensions, which would be less able to establish functional connections.

In the co-cultures, we observed a significant increase on cell death in both neural and muscle populations upon MN-restricted *FUS* mutation or HuD overexpression. Both *in vivo* and *in vitro*, MNs are recognized for their ability to release trophic factors that support the survival and growth of nearby cells. The presence of MNs in the co-culture can thus enhance the survival of skeletal muscle cells and contribute to their overall health. We hypothesize that upon increased FUS^P525L^ and FUS^WT^+HuD MNs death and/or failure to form mature NMJs, their support to muscle survival in vitro fails, thus explaining the increased apoptosis of muscle fibers.

The case of the sporadic ALS patient who died unexpectedly (Fischer et al., 2004), despite anecdotal, suggests that at the time of diagnosis (first signs of motor symptoms) MN cell bodies might still be intact. Thus, early intervention aimed to restore NMJs might result in an effective therapeutic approach to prevent irreversible subsequent MN degeneration. To this regard, our work shows that individual knockdown of HuD and one of its key targets, NRN1, can rescue endplate formation and apoptosis in vitro. Moreover, *in vivo* evidence from the *Drosophila* model shows that reduction of the *HuD*-related gene *elav* expression is sufficient to rescue the motor phenotype caused by FUS completely. Here we took advantage of RNA interference. However, recent advances in the development of antisense oligonucleotides (ASOs) for therapy, also in the context of ALS (McCampbell et al., 2018), provide the opportunity to translate these findings to patients in the near future.

## MATERIALS AND METHODS

### iPSC culture, transfection and differentiation

Human iPSC lines were maintained in Nutristem-XF medium (Biological Industries), with 0.1X Penicillin/Streptomycin (Merck Life Sciences) (Nutristem P/S), in Matrigel-coated dishes (hESC-qualified Matrigel; Corning) and passaged every 4-5 days with 1 mg/ml Dispase (Gibco). The human FUS^WT^ line, its isogenic FUS^P525L^ line, and the FUS^WT^+HuD line had been previously derived from the WT-I line (Lenzi et al., 2015; Garone et al., 2021). The FUS^WT^ iPSCs line WTSIi004-A (WTSI-FUS^WT^) was obtained by the European Bank for induced pluripotent Stem Cells (EBiSC) and used to generate an independent set of iPSC lines, considered here as a biological replicate of the WT-I-based set of iPSC lines. From these cells we generated the isogenic WTSI-FUS^P525L^ iPSC line by TALEN-directed mutagenesis as previously described (Lenzi et al., 2015). Both mutant and control lines went through the gene editing process. To generate the iPSC line WTSI-FUS^WT^ overexpressing HuD (WTSI-FUS^WT^+HuD), cells were co-transfected with 4.5 µg of transposable vector epB-Puro-TT-SYN1-HuD (SYN1::HuD) and 0.5 µg of the piggyBac transposase using the Neon Transfection System (Life Technologies) as described (Garone et al., 2021). Stable and inducible iPSCs for spinal MN differentiation (NIL iPSCs) were obtained as previously described (De Santis et al., 2018; Garone et al., 2019). Briefly, iPSCs were transfected with 4.5 μg of NIL piggyBac construct and 0.5 μg of transposase vector using the Neon Tranfection System (Life Technologies) and then selected with 5 μg/ml blasticidin in Nutristem-XF medium containing 0.1X P/S for 10 days. KOLF-1 (FUS^WT^) iPSCs, stably transfected with inducible transposable vectors encoding for BAF60c and MyoD1 (Tiago et al., 2021) (MB iPSCs), were used to generate SkMCs. Briefly, this line was generated by co-transfecting iPSCs with 2.25 μg each of the ePB-Bsd-TT-BAF60c and ePB-Puro-TT-m-MyoD vectors, and 0.5 μg of piggyBac transposase, and then selected with 1 μg/ml puromycin and 5μg/ml blasticidin in Nutristem P/S for 10 days, as previously described (Lenzi et al., 2016)

### Differentiation of spinal MNs and SkMCs and co-cultures

Spinal MNs were obtained as previously described (De Santis et al., 2018; Garone et al., 2019). Briefly, NIL iPSCs were dissociated with Accutase (Thermo Fisher Scientific) to single cells and replated in Nutristem P/S supplemented with 10 µM Y-27632 ROCK inhibitor (Enzo Life Sciences). The next day differentiation was induced in DMEM/F12 (Dulbecco’s Modified Eagle’s Medium/ Nutrient Mixture F-12 Ham; Merck Life Sciences), 1X Glutamax (Thermo Fisher Scientific), 1X NEAA (Thermo Fisher Scientific), 0.5X P/S and doxycycline 1 µg/ml (Thermo Fisher Scientific) for 2 days. On the third day, medium was replaced by Neurobasal/B27 (Neurobasal medium, Thermo Fisher Scientific; 1X B27, Thermo Fisher Scientific; 1X Glutamax; 1X NEAA; 0.5X P/S) supplemented with 5 µM DAPT, 4 µM SU5402 (both from Merck Life Sciences) and 1 µg/ml doxycycline for 3 days. At day 5, MN progenitors were dissociated with Accutase and counted. 1×10^6^ cells were plated onto Matrigel-coated 35 mm dishes for RNA analyses, and 5×10^4^ cells per well were plated onto Matrigel-coated μ-Slide 8 Well (ibidi) for immunofluorescence analyses. 10 μM of Y-27632 ROCK inhibitor was added for 24h after dissociation. Neurons were maintained in Neurobasal/B27 supplemented with 20 ng/ml L-ascorbic acid (Merck Life Sciences), 20 ng/ml BDNF (PreproTech) and 10 ng/ml GDNF (PreproTech).

To obtain SkMCs, we adapted a method to convert iPSCs into SkMCs upon ectopic expression of MyoD and BAF60c (Lenzi et al., 2016; Caputo et al., 2020). This protocol has been previously developed in our lab for short-term cultures (up to 9 days), however we noticed that a subpopulation of differentiation resistant cells persisted in longer cultures. Over time, these mitotically active cells outnumbered terminally differentiated SkMCs, thus hindering the establishment of long-term co-cultures. To overcome this problem, we included a step consisting of the transient addition of cytosine arabinoside (AraC), which successfully reduced the number of proliferating undifferentiated cells, as follows. MB iPSCs were dissociated with Accutase (Thermo Fisher Scientific) to single cells and replated in Nutristem P/S supplemented with 10 µM Y-27632 in 35 mm dishes (2×10^5^ cells). In the following two days, cells were maintained with Growth Medium consisting of high glucose DMEM (Merck Life Sciences), 1X Glutamax, 1X P/S, 20% FBS (Merck Life Sciences), 50 μg/ml Insulin (Roche), 25 ng/ml bFGF (Corning) and 10 ng/ml hEGF (Corning) and 200 ng/ml doxycycline. At day 2, medium was replaced by Skeletal Muscle Differentiation Medium (Promega), 1X P/S, 200 ng/ml doxycycline and 2μM AraC (Merck Life Sciences). At day 4, this medium was replaced by Skeletal Muscle Differentiation Medium, 1X P/S and 200 ng/ml doxycycline. The medium was then changed every other day.

To generate co-cultures of spinal MNs and SkMCs we adapted previously published protocols (Guo et al., 2011; Stoklund Dittlau et al., 2021). MB iPSCs were dissociated and replated in 35 mm dishes (2×10^5^ cells) or μ-Slide 8 Well (ibidi; 2×10^4^ cells per well) and then SkMC differentiation was induced as described above. Spinal MNs differentiation started independently and in parallel when SkMCs were at day 7 of the muscle differentiation protocol. At day 5, MNs progenitors were replated (at approximately 1:1 ratio) in the 35mm dishes (5×10^5^ cells) or μ-Slide 8 Well (ibidi; 5×10^4^ cells per well) containing SkMCs, which at this stage had reached day 13 of differentiation, in Neurobasal/B27 medium containing 20 ng/ml L-ascorbic acid, 20 ng/ml BDNF, 10 ng/ml GDNF, 200 ng/ml doxycycline and 10 µM Y-27632. The next day, medium was replaced with Neurobasal/B27, 20 ng/ml L-ascorbic acid, 20 ng/ml BDNF, 10 ng/ml GDNF, 200 ng/ml doxycycline, 20 μg/ml laminin (Merck Life Sciences) and 0.01 μg/ml agrin (R&D system). Co-cultures were maintained for 14, 28 or 50 days, changing medium every 4 days.

### Functional assays in iPSC-derived cells

Functional activity in co-cultures and muscle monocultures was assessed by adding 50μM glutamate (Gibco) to cells previously plated in μ-Slide 8 Well (ibidi) supports and maintained in Neurobasal/B27 medium containing 20 ng/ml L-ascorbic acid, 20 ng/ml BDNF and 10 ng/ml GDNF. Five minutes after the addition of glutamate, the number of muscle contractions per minute in 3 randomly selected fields was counted under Carl Zeiss Axio Vert.A1 Microscope. To verify muscle contraction inhibition, 10μM tubocurarine (Sigma-Aldrich) was added to the medium containing glutamate. Five minutes after the addition of tubocurarine, the same co-culture fields were identified for measuring muscle contractions per minute. These experiments were performed in 3 independent batches of differentiated iPSCs.

### Real-Time qRT-PCR

Total RNA was extracted using E.Z.N.A Total RNA Kit I (Omega BIO-TEK) and retrotranscribed with iScript Supermix (Bio-Rad Laboratories) according to manufacturers’ instructions. qRT-PCR was performed with the iTaq Universal SYBR Green Supermix kit (Bio-Rad Laboratories) on a ViiA 7 system (Thermo Fisher Scientific). *ATP5O* and *TUBB3* were employed as calibrator genes. Full list of primers is reported in Supplementary Table S1.

### Cell immunofluorescence and confocal imaging

Cells were fixed in 4 % PFA (Electron microscopy sciences) and 30 mM sucrose (Sigma-Aldrich) for 10 min at room temperature (RT), then washed with PBS with calcium chloride and magnesium chloride (Merck Life Sciences) and permeabilized with a solution of 0.5% BSA, 10% Horse Serum (HRS), 0.2% Triton X-100 and PBS (Merck Life Sciences) for 15 min. Cells were then washed and treated with a blocking solution composed of 0.5% BSA, 10% HRS and PBS for 30 min. For NMJ staining, anti-MyHC (1:50; MAB4470, R&D system) and anti-SYP (1:200; ab32127, Abcam) were diluted in blocking solution and incubated at 4°C overnight. Anti-TUJ1 (1:1000; T2200, Merck Life Sciences) was diluted in the same solution and incubated for 1 h at RT. Cells were then washed with PBS and incubated with α-bungarotoxin-488 (1:350, B13422, Thermo Fisher Scientific) for 1 h at RT. Secondary anti-mouse Alexa Fluor 647 (1:250, Immunological Science) and anti-rabbit Alexa Fluor 594 (1:250, Immunological Sciences) antibodies were then incubated for 1 h. DAPI (1:2000; Sigma-Aldrich) diluted in 0.5% BSA, 10% HRS in PBS was added for 5 minutes. Cells were washed in PBS and ibidi Mounting Medium (50001, ibidi) was added. Confocal image stacks were acquired using an inverted Olympus iX73 equipped with an X-light V3 spinning disc head (Crest Optics), a LDI laser illuminator (89 North), Prime BSI sCMOS camera (Photometrics) and MetaMorph software (Molecular devices) with 40X air or 100X oil objectives (Olympus). For skeletal muscle with or without 48 h Ara-C treatment, fixed cells were incubated with anti-MyHC (1:50; MAB4470, R&D system) and anti-NANOG (1:200; PA1-097X; Thermo Fisher Scientific) primary antibodies overnight. Then, anti-mouse Alexa Fluor 647 (1:250, Immunological Science), anti-rabbit Alexa Fluor 488 (1:250, Immunological Science) and DAPI were used and mounted with ibidi Mounting Medium (50001, ibidi). Confocal image stacks were acquired using an Olympus FV1200 confocal laser scanning microscope with 20X air-objective. For α-BTX positive fibers analysis, randomly selected fields for each batch of differentiated iPSCs, considered as a replicate, were used to count the total number of muscle fibers (based on the MyHC staining), and the total number of muscle fibers showing α-BTX signal to calculate the percentage. For TUJ1 analysis, we calculated the average value of randomly selected fields for each replicate.

### Cell-based assays and confocal imaging

For apoptosis analysis, fixed cells were incubated with anti-CC-3 (1:400, 9661, Cell signaling) overnight, anti-MAP2 (1:2000, ab5392, Abcam) for 1h at RT. Anti-chicken Alexa Fluor 488 (1:250, Immunological Sciences) and anti-rabbit Alexa Fluor 594 (1:250, Immunological Sciences) secondary antibodies were then used. DAPI was added for 5 minutes. Confocal image stacks were acquired using an Olympus FV10i laser scanning confocal microscope with 10X air objective at 2x zoom or the X-light V3 spinning disc confocal microscope described before with 20X air objective (Olympus). Phase-contrast images were acquired using Carl Zeiss Axio Vert.A1 Microscope with 20X or 40X air objectives. For dead cell quantification, live co-cultures were incubated (5% CO_2_ at 37°C) with Neurobasal/B27, 20 ng/ml L-ascorbic acid, 20 ng/ml BDNF and 10 ng/ml GDNF medium supplemented with 0.6 μl ethidium homodimer-1 594 (Thermo Fisher Scientific) according to manufacturer’s protocol. After 30 minutes cells were washed with PBS and maintained in Neurobasal/B27, 20 ng/ml L-ascorbic acid, 20 ng/ml BDNF and 10 ng/ml GDNF medium during acquisition. Confocal image stacks were acquired using Olympus FV10i laser scanning confocal microscope with 10x air objective at 1.5x zoom.

### RNA interference (RNAi) in iPSC-derived in co-cultures

Lyophilized siRNAs were resuspended in nuclease-free water at 20μM stock concentration. HuD, NRN1, GAP-43 and non-targeting siRNAs (siRNA-SMARTpool and ON-TARGET plus Non-targetingPool, Dharmacon) were transfected at 10 nM concentration using siLentFect Lipid Reagent (Bio-Rad Laboratories) according to manufacturer’s instructions. Medium was replaced 5 h after transfection.

### Drosophila melanogaster stocks

All *Drosophila* stocks were maintained on standard cornmeal medium at 25°C in light/dark-controlled incubators. The UAS-FUS WT and UAS-FUS P525L lines were generated through site specific insertion of the transgene at BestGene Inc. using the (attp2) integration vector and were previously described (Casci et al., 2019; Anderson et al., 2018). The TRIP control lines, the UAS-Luciferase (*luc* OE, # 35789) line, and the UAS-elav (*elav* OE, stock # 38400) line were obtained from Bloomington Drosophila Resource Center (BDSC). The UAS-elav RNAi (#37915) was obtained from the Vienna Drosophila Resource Center (VDRC). The Elav-GeneSwitch-Gal4 is a gift from Dr. Haig Keshishian at Yale University.

### Drosophila melanogaster motor function assays

The motor function phenotypes of all control and diseased conditions were collected at 25°C. Elav-GS-Gal4 (neuron specific driver) was used to express FUS-WT and FUS-P525L with and without knockdown of ELAV. For performing the climbing assay, flies were crossed in the absence of RU486 (mifepristone, Cayman Chemical, #10006317) on standard food. Day 1 adult females were transferred to RU486 (5mM) drug food and placed at 25°C for 10-15 days before performing the motor function assay at the time points indicated. 6-10 flies were transferred to empty rounded vials and allowed to acclimate for 20 minutes in the new vials. Flies were knocked to the bottom of the vials by tapping each vial against the laboratory bench three times, and a video camera was used to record flies climbing up the walls. The percentage of flies that climbed 4cm in 10s and the velocity of each fly was quantified and analyzed using GraphPad PRISM 8.0 software. Three experimental replicates (trials) were performed for each group.

### Drosophila melanogaster NMJ analysis

All the genetic crosses were crossed with the inducible pan-neuronal driver, Elav-GS-Gal4, at 25°C on standard media fed with 5µM final concentration of RU-486 drug (#10006317, Cayman Chemicals). Third-instar larvae were collected for NMJ analysis. For immunofluorescence, the larvae were rinsed with PBS and dissected along the dorsal midline to expose the NMJs. Dissected Drosophila tissues were washed with PBS (Lonza) and fixed with 4% paraformaldehyde (Sigma-Aldrich, P6148) for 20 minutes at room temperature. The tissues were washed three times with PBS following fixation and blocked with blocking buffer: 5% normal goat serum (NGS, Abcam, AB7681) in PBS with 0.1% Triton-X (PBST). The samples were incubated with primary antibodies overnight at 4°C and washed four times with 0.1% PBST, incubated with secondary antibodies for 2h at room temperature followed by four washes with PBST. Samples were mounted with Fluoroshield (Sigma-Aldrich, F6057) onto glass-slides. Primary and secondary antibodies were prepared in a blocking buffer. Primary antibodies: Cy3 conjugated Horse Radish Peroxidase (1:200; Jackson ImmunoResearch, 123-165-021), mouse Cysteine String Protein (dCSP) (1:100; Developmental Studies Hybridoma Bank). Secondary antibodies: Goat anti-mouse Alexa Fluor 488 (1:1000; Invitrogen, A11008); Goat anti-rabbit Alexa Fluor-568 (1:100; Invitrogen, #651727). Confocal images were acquired in Z-stacks using a Nikon A1-T216.3 confocal microscope at 60X (oil) magnification. NMJs innervating Muscle 4 at hemi-segments A3-A4 were used for analyzing synaptic bouton quantities. For each NMJ, the number of synaptic boutons per NMJ was normalized to the surface area of the innervating muscle. The bouton number and the NMJ area were quantified using ImageJ (NIH).

### Oxidative stress induction and transcription block

Cells were left untreated or treated with 0.5 mM sodium arsenite (Merck Life Sciences) to induce acute oxidative stress for 1h. For transcription inhibition, 25 μl/ml α-amanitin solution (Merck Life Sciences) was added to the medium for 8h; in the last hour, 0.5 mM sodium arsenite was added to induce oxidative stress.

### Bioinformatics analysis of sporadic ALS RNA-seq datasets

The raw count matrix relative to the RNA-seq analysis of ALS patient samples and individuals with and without other neurological disorders produced by the NYGC consortium (Tam et al., 2019) was retrieved from the Gene Expression Omnibus (GEO) database (Edgar et al., 2002) (accession number: GSE124439), while the metadata information was taken from the original publication. Starting from this matrix, sample-specific gene-level FPKM (Fragments Per Kilobase of exon per Million mapped reads, a normalized expression measure) values were calculated using the “fpkm” function from the DESeq2 v1.32.0 R package (Love et al., 2014) and the exonic gene size information retrieved from the GENCODE v42 annotation (Harrow et al., 2012). Differential expression analysis between different conditions was performed using the “DESeq” function, controlling for sex and tissue type.

### Statistics and reproducibility

Statistical analysis, graphs, and plots were generated using GraphPad PRISM 8.0 (GraphPad Software). The statistical test used is indicated in each figure legend. Sample size and the definition of replicates for each experiment is also indicated in the figure legends. We deemed statistically significant *p* values at a threshold of less than 0.05. The adjusted *p* values obtained from differential expression analyses shown in figure Figure 7G and Supplementary Figure S8A have been corrected for multiple comparisons via the Benjamini-Hochberg procedure.

## DATA AVAILABILITY

All data generated or analyzed during this study are included in the manuscript and supporting files. Materials are available upon request.

## AUTHOR CONTRIBUTION

Conceptualization, B.S., A.R.; Formal analysis, B.S., M.M., D.M., A.C.; Investigation, B.S., D.M.; Methodology, B.S., M.M., A.C., B.B., V.d.T., M.M., C.P.Z., M.G.G.; Project administration, A.R.; Supervision, U.B.P., A.R.; Writing - original draft, A.R.; Writing - Review & Editing, B.S., U.B.P., A.R.. All authors read and approved the final manuscript.

## ACKNOWLEDGEMENTS

The authors wish to thank the Imaging Facility at Center for Life Nano and Neuro-Science, Fondazione Istituto Italiano di Tecnologia, for support and technical advice. We thank Dr. Tiziana Santini and Dr. Ersilia Fornetti for advice on NMJ staining and analysis. We thank Dr. R. De Santis, Prof. I. Bozzoni, prof. S. Di Angelantonio and the members of the Rosa lab for helpful discussion.

Project funded under the National Recovery and Resilience Plan (NRRP), project “National Center for Gene Therapy and Drug based on RNA Technology” (CN00000041). Financed by NextGenerationEU PNRR MUR – M4C2 – Action 1.4-Call “Potenziamento strutture di ricerca e di campioni nazionali di R&S” (CUP: B83C22002870006) to AR. This work has been supported by National Institute of Health (NIH) grant R01NS081303 to U.B.P.

