## Supplementary material for "HuD (ELAVL4) gain-of-function impairs neuromuscular junctions and induces apoptosis in *in vitro* and *in vivo* models of amyotrophic lateral sclerosis"

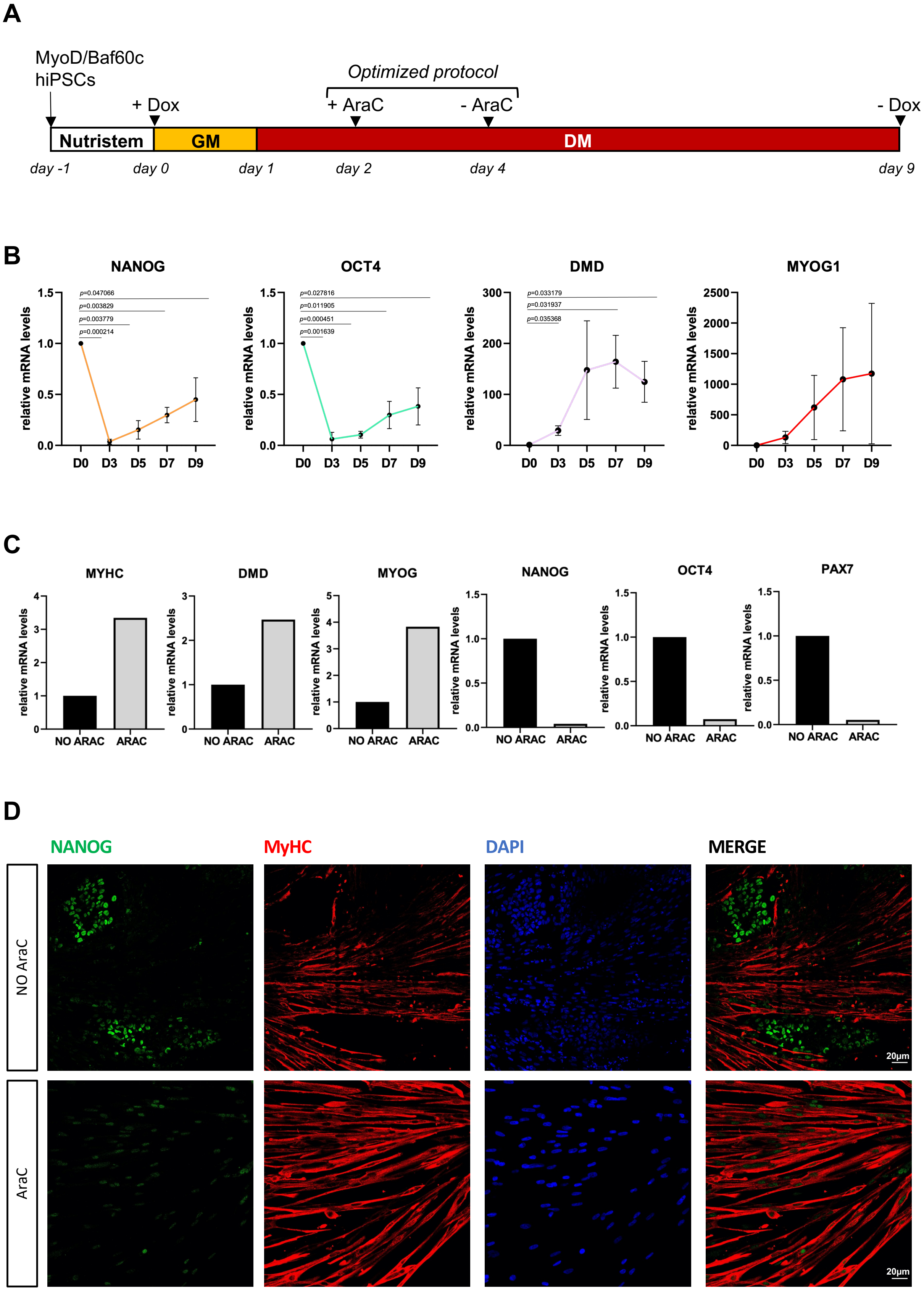
A
MyoD/Baf60c
hiPSCs
+ Dox
Optimized protocol
+ AraC
- AraC
- Dox
Nutristem
GM
DM
day -1
day 0
day 1
day 2
day 4
day 9
B
NANOG
OCT4
DMD
MYOG1
relative mRNA levels
p=0.047066
p=0.003829
p=0.003779
p=0.000214
p=0.027816
p=0.011905
p=0.000451
p=0.001639
p=0.033179
p=0.031937
p=0.035368
D0
D3
D5
D7
D9
C
MYHC
DMD
MYOG
NANOG
OCT4
PAX7
relative mRNA levels
NO ARAC
ARAC
D
NANOG
MyHC
DAPI
MERGE
NO AraC
AraC
20μm
20μm

**Supplementary Figure S1. Optimization of the skeletal muscle differentiation protocol. Related to Figure 1.**

(A) Schematic representation of the iPSC differentiation method to obtain SkMCs. (B) qRT-PCR expression analysis of pluripotency (*NANOG*, *OCT4*) and differentiation (*DMD*, *MYOG1*) markers, normalized on ATP5O expression, at the indicated time points (Day 0, 3, 5, 7, 9) from differentiation experiments performed without the use of cytosine arabinoside (AraC) at days 2 to 4 of the protocol, i.e. using the original method described in (Lenzi et al., 2016). Error bars indicate standard deviation (n=3 Student's t-test, paired, two tails). (C) Comparative qRT-PCR analysis of the expression of stemness/pluripotency (*PAX7*, *OCT4*, *NANOG*) and muscle (*DMD*, *MYOG1* and *MYHC*) markers at day 13 of differentiation (n=1), using the original (NO AraC) or optimized (AraC) protocol. ATP5O expression was used for normalization. (D) Immunofluorescence analysis of SkMCs as in (C). Nuclei were stained with DAPI (blue). Scale bar: 20μm.

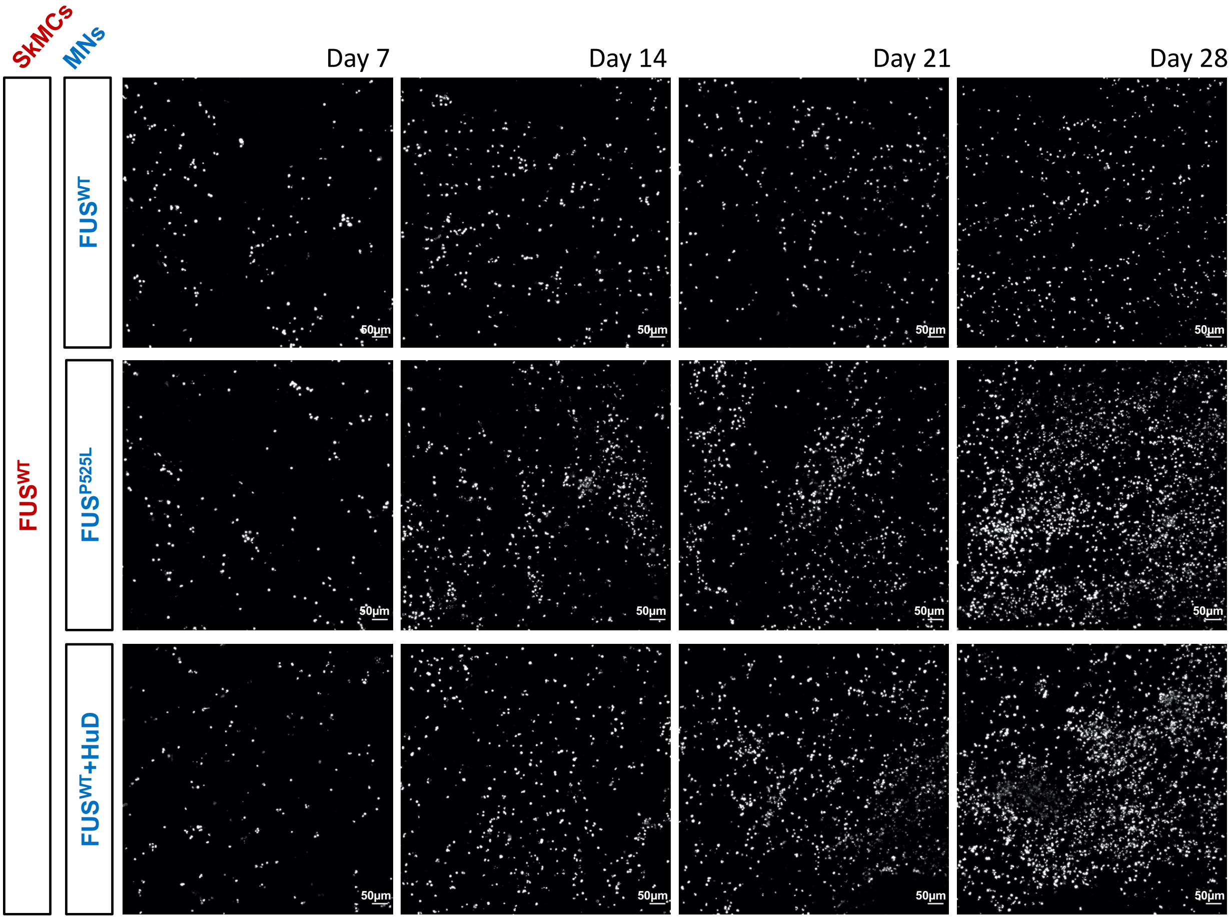


**Supplementary Figure S2. Dead cell staining. Related to Figure 4.**
Representative images of co-cultures stained with Ethidium Homodimer-1 (gray), used for quantification shown in Figure 4B. Scale bar: 50µm.

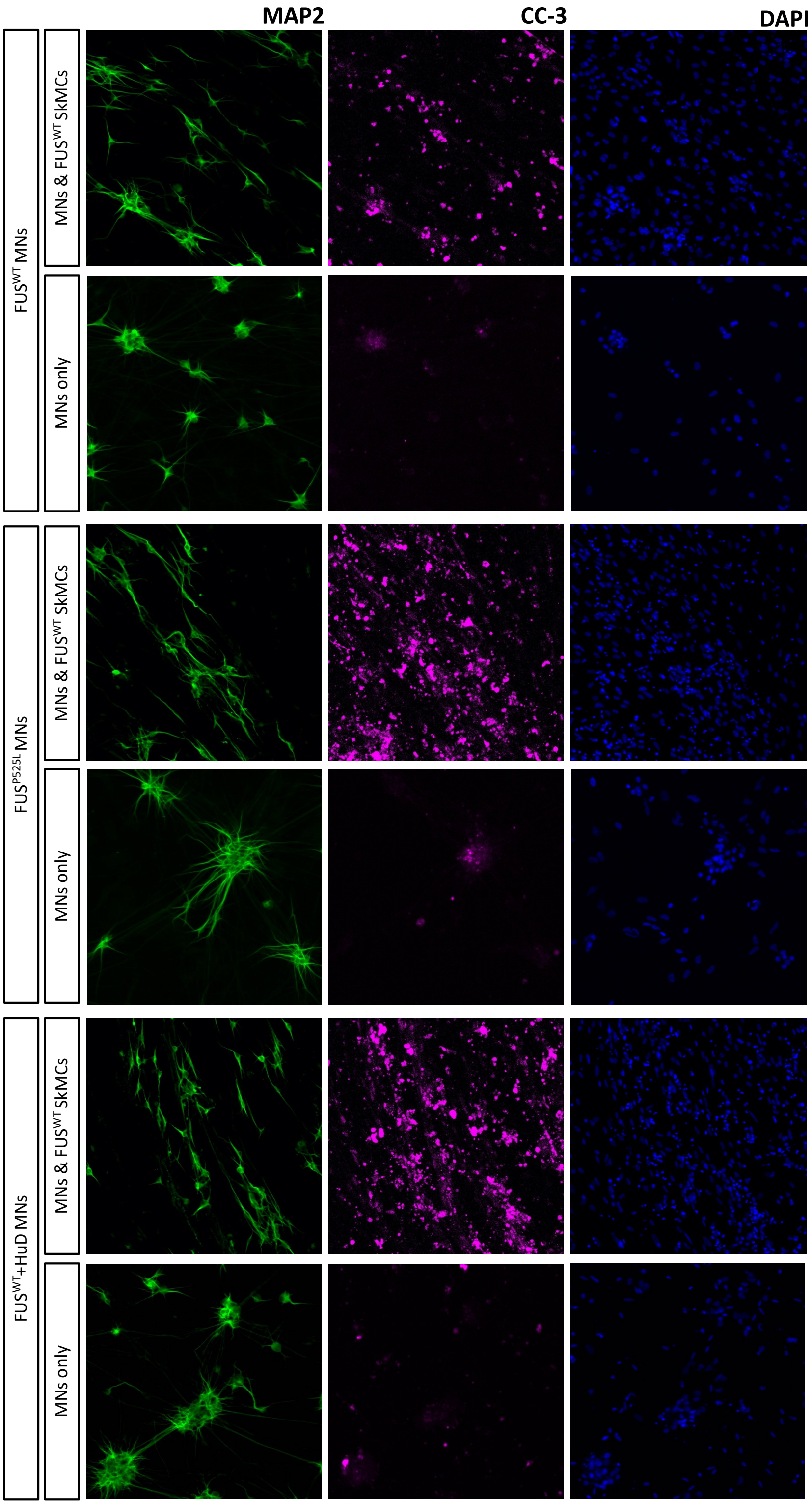


**Supplementary Figure S3. Single panels of immunofluorescence staining. Related to Figure 4.**

Single panels of immunofluorescence staining, shown in Figure 4C as merged images, of co-cultures or MN monocultures at day 14, using Cleaved Caspase-3 (CC-3) and MAP2 antibodies, and DAPI.

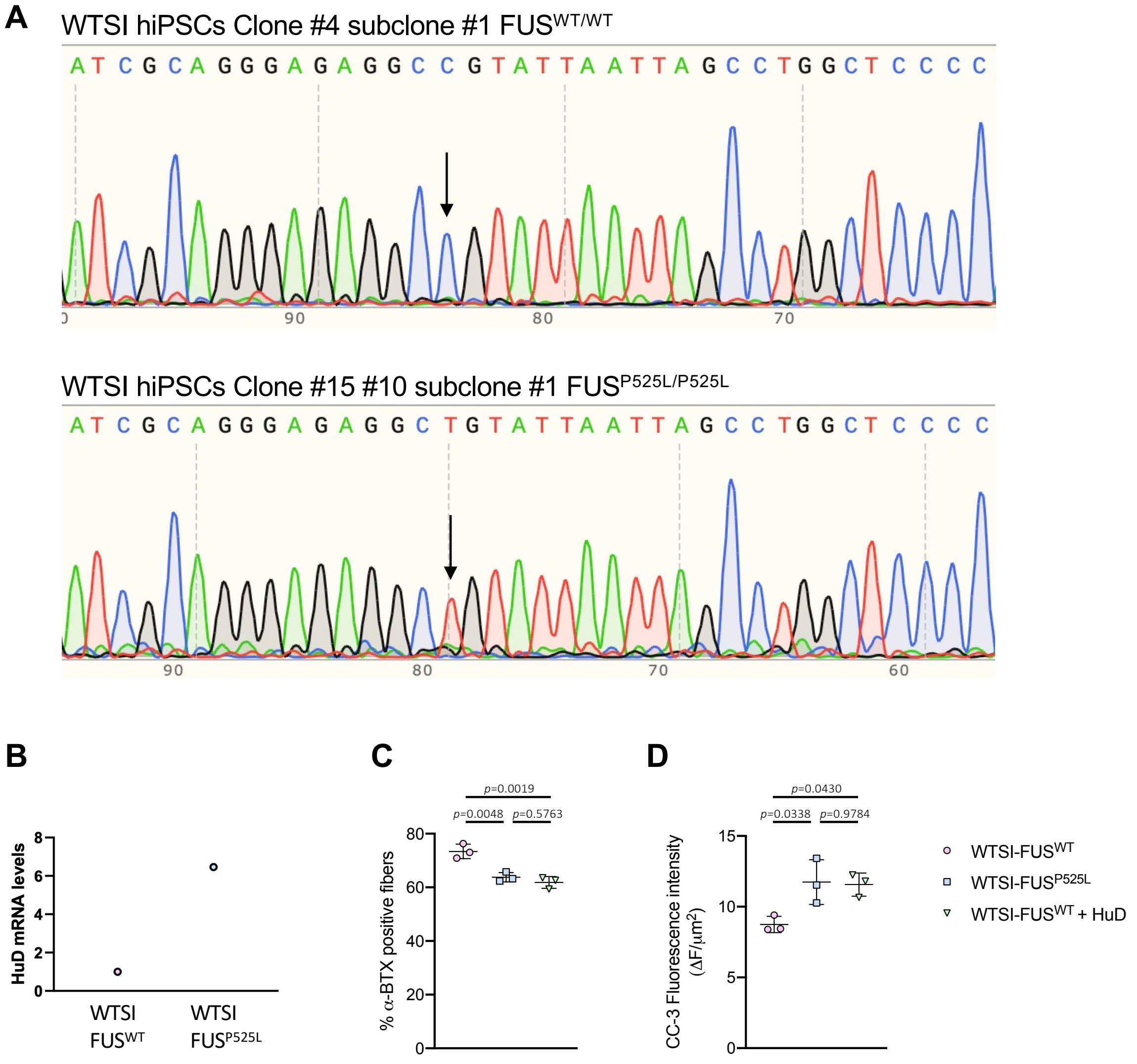


**Supplementary Figure S4. Generation and characterization of the new FUS$^{P525L}$ mutant line derived from WTSIi004-A iPSCs. Related to Figure 4.**

(A) Sanger sequencing of a wild-type clone (top) and a FUS$^{P525L}$ mutant (bottom) clone derived from the WTSIi004-A iPSC line (EBiSC) as previously described (Lenzi et al., 2015). These lines are named WTSI-FUS$^{WT}$ and WTSI-FUS$^{P525L}$ in the present paper. Arrows indicate the targeted nucleotide in codon 525: C, wild type; T, mutant. (B) Analysis of *HuD* mRNA levels in WTSI-FUS$^{WT}$ and WTSI-FUS$^{P525L}$ spinal MNs at day 12 of differentiation. TUBB3 has been used for normalization (n=1). (C,D) The graphs report quantitative analysis of co-cultures made with FUS$^{WT}$ SkMCs and WTSI-FUS$^{WT}$, WTSI-FUS$^{P525L}$, or WTSI-FUS$^{WT}$+HuD spinal MNs at day 14: percentage of α-BTX positive fibers (C) and CC-3 fluorescence intensity (D). Each dot represents a replicate, consisting of an individual batch of differentiated iPSCs. For each replicate, 6 fields have been acquired for α-BTX positive fibers analysis, and 8-10 for CC-3 immunofluorescence. The dot shows the average value. Error bars indicate standard deviation calculated on the average value of the replicates. Ordinary one-way ANOVA, post hoc Tukey test $p$ values for multiple comparisons are indicated in the graph.

A

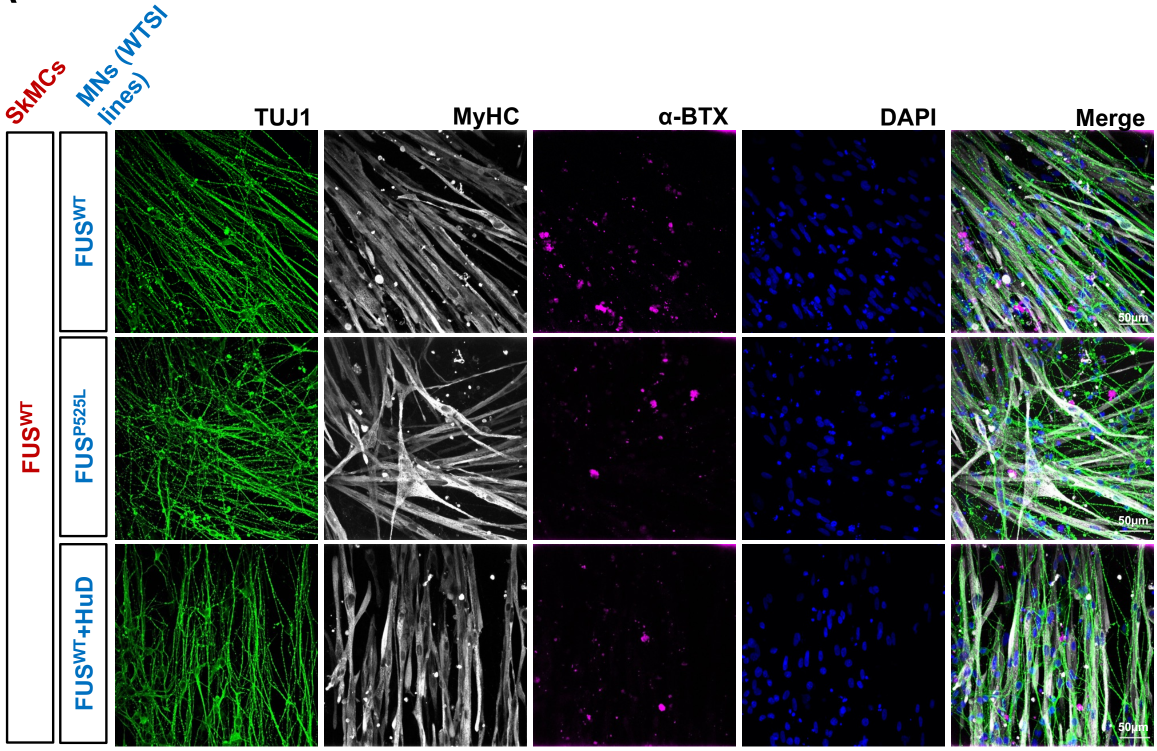


B

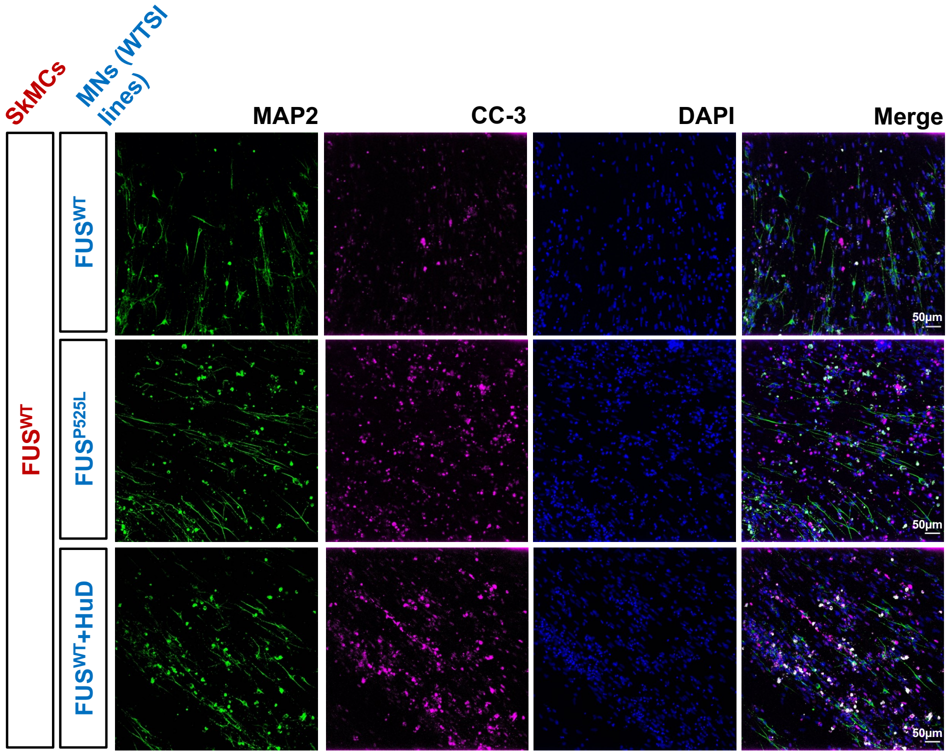


**Supplementary Figure S5. Images used for quantification (WTSI co-cultures). Related to Figures 2 and 4.**

(A-B) Images used for quantification analyses of Supplementary Figure S4C (panel A) and Supplementary Figure S4D (panel B). Scale bar: 50µm.

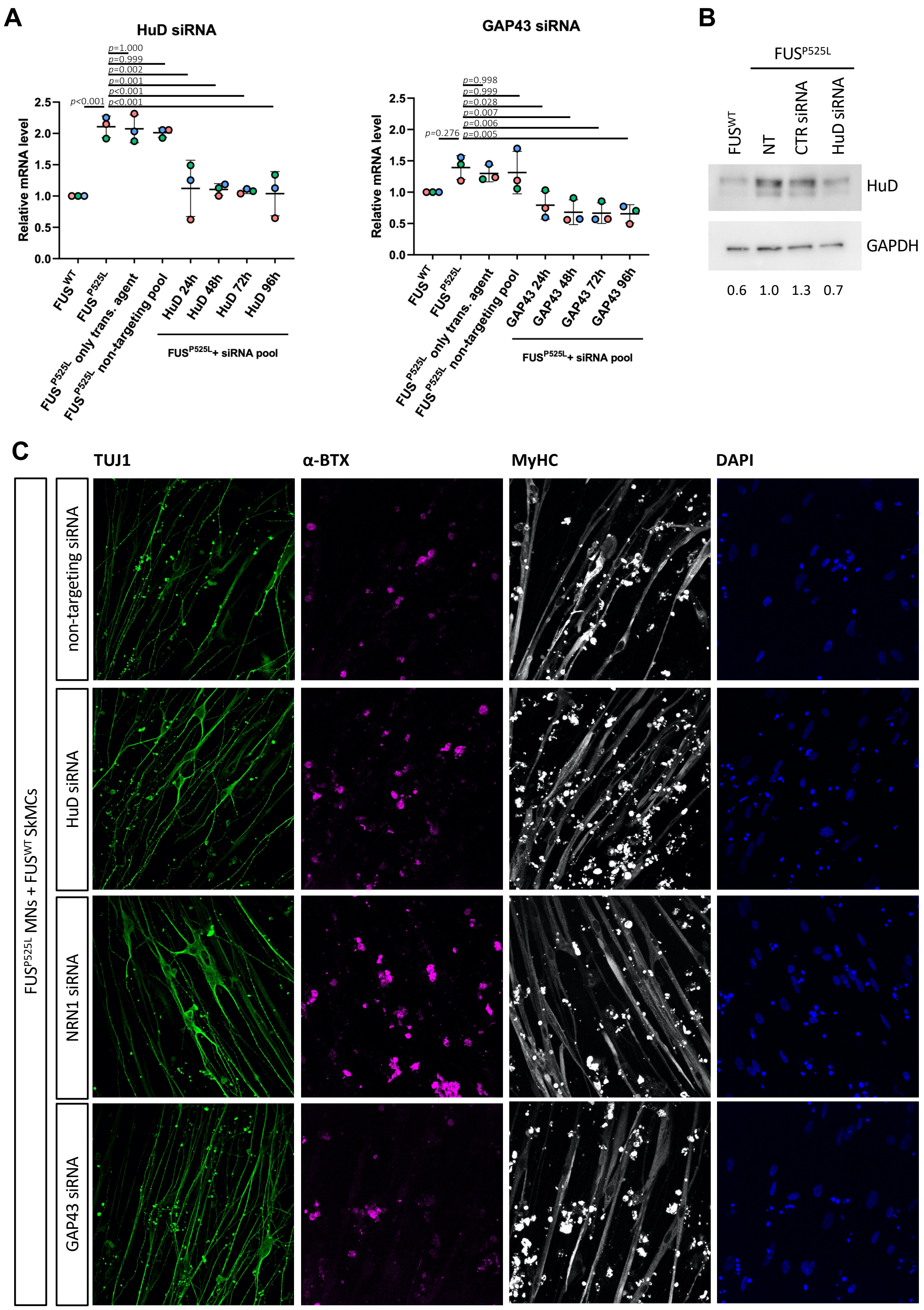
A
HuD siRNA
Relative mRNA level
p<0.001
p=1.000
p=0.999
p=0.002
p=0.001
p<0.001
p<0.001
FUSWT
FUSP525L
FUSP525L only trans. agent
FUSP525L non-targeting pool
HuD 24h
HuD 48h
HuD 72h
HuD 96h
FUSP525L+ siRNA pool
GAP43 siRNA
p=0.276
p=0.998
p=0.999
p=0.028
p=0.007
p=0.006
p=0.005
GAP43 24h
GAP43 48h
GAP43 72h
GAP43 96h
B
FUSP525L
FUSWT
NT
CTR siRNA
HuD siRNA
HuD
GAPDH
0.6
1.0
1.3
0.7
C
TUJ1
α-BTX
MyHC
DAPI
FUSP525L MNs + FUSWT SkMCs
non-targeting siRNA
HuD siRNA
NRN1 siRNA
GAP43 siRNA

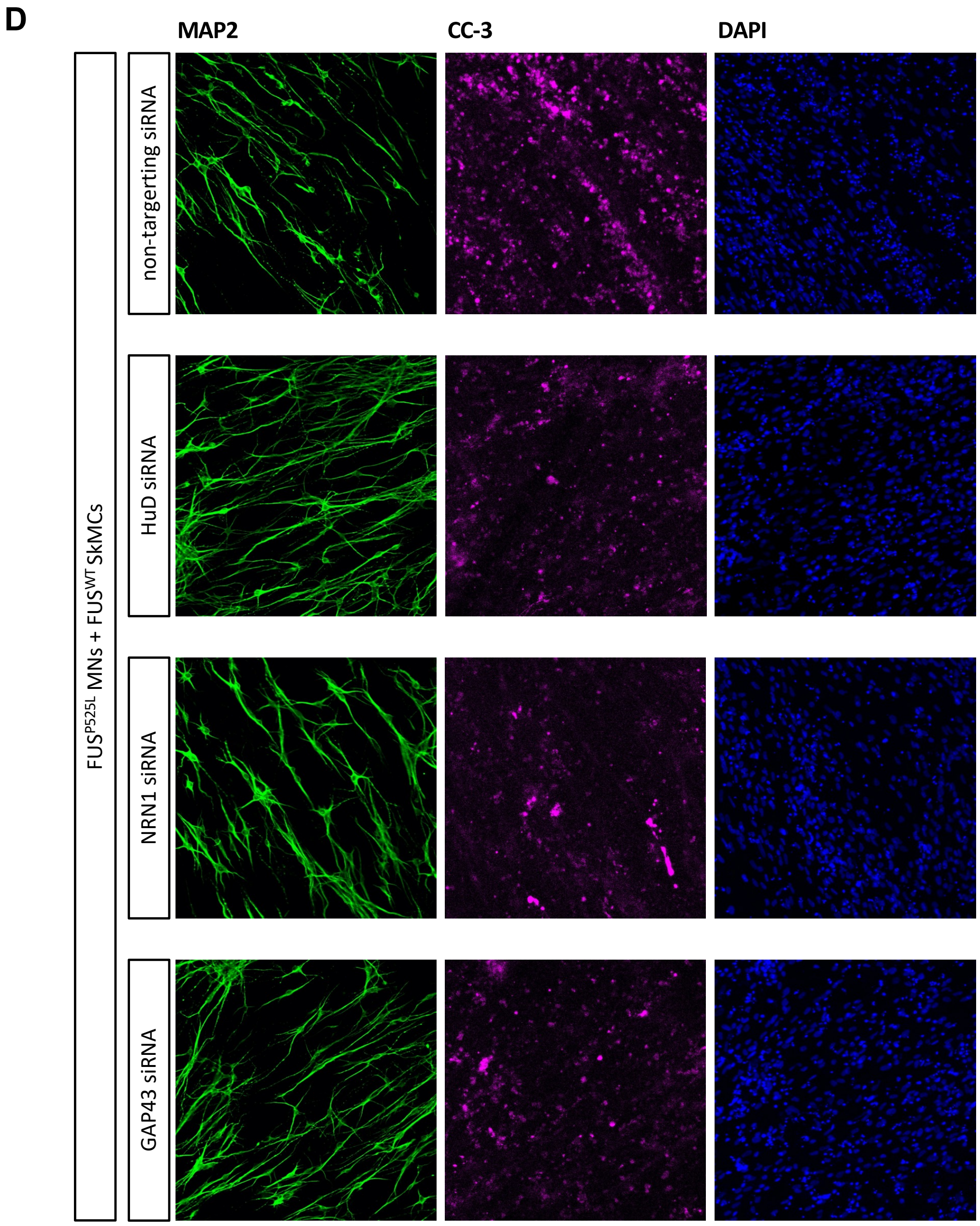


**Supplementary Figure S6. Knockdown by RNAi in FUS$^{P525L}$ MNs and co-cultures. Related to Figure 5.**

(A) Analysis of *HuD* (left) and *GAP43* (right) mRNA levels in untransfected FUS$^{WT}$ and FUS$^{P525L}$ spinal MNs and FUS$^{P525L}$ spinal MNs transfected with non-targeting control siRNA or siRNAs targeting *HuD* or *GAP43*. *ATP5O* has been used for normalization. Values indicated in the graphs are relative to untransfected FUS$^{WT}$ samples. Each dot represents a replicate, consisting of an individual batch of differentiated iPSCs. The graphs show the average and standard deviation. Ordinary one-way ANOVA, post hoc Tukey test *p* values for multiple comparisons of all other conditions with the FUS$^{P525L}$ untransfected sample are indicated in the graph. The same analysis for validating siRNAs targeting NRN1, used in the present work, had been previously reported (Garone et al., 2021). (B) Western blot analysis of HuD protein levels in FUS$^{WT}$ and FUS$^{P525L}$ spinal MNs (NT) and FUS$^{P525L}$ spinal MNs transfected for 48 hrs with non-targeting control siRNA (CTR siRNA) or siRNAs targeting *HuD* (HuD siRNA). GAPDH was used as a loading control. The numbers indicate the quantification of HuD signals normalized for GAPDH signals, relative to the FUS$^{P525L}$ NT sample set as 1. Uncropped images are shown in Supplementary Figure S11. (C,D) Single panels of the immunofluorescence staining, shown in Figure 5A,B as merged images, of co-cultures made with $FUS^{WT}$ SkMCs and $FUS^{P525L}$ MNs.

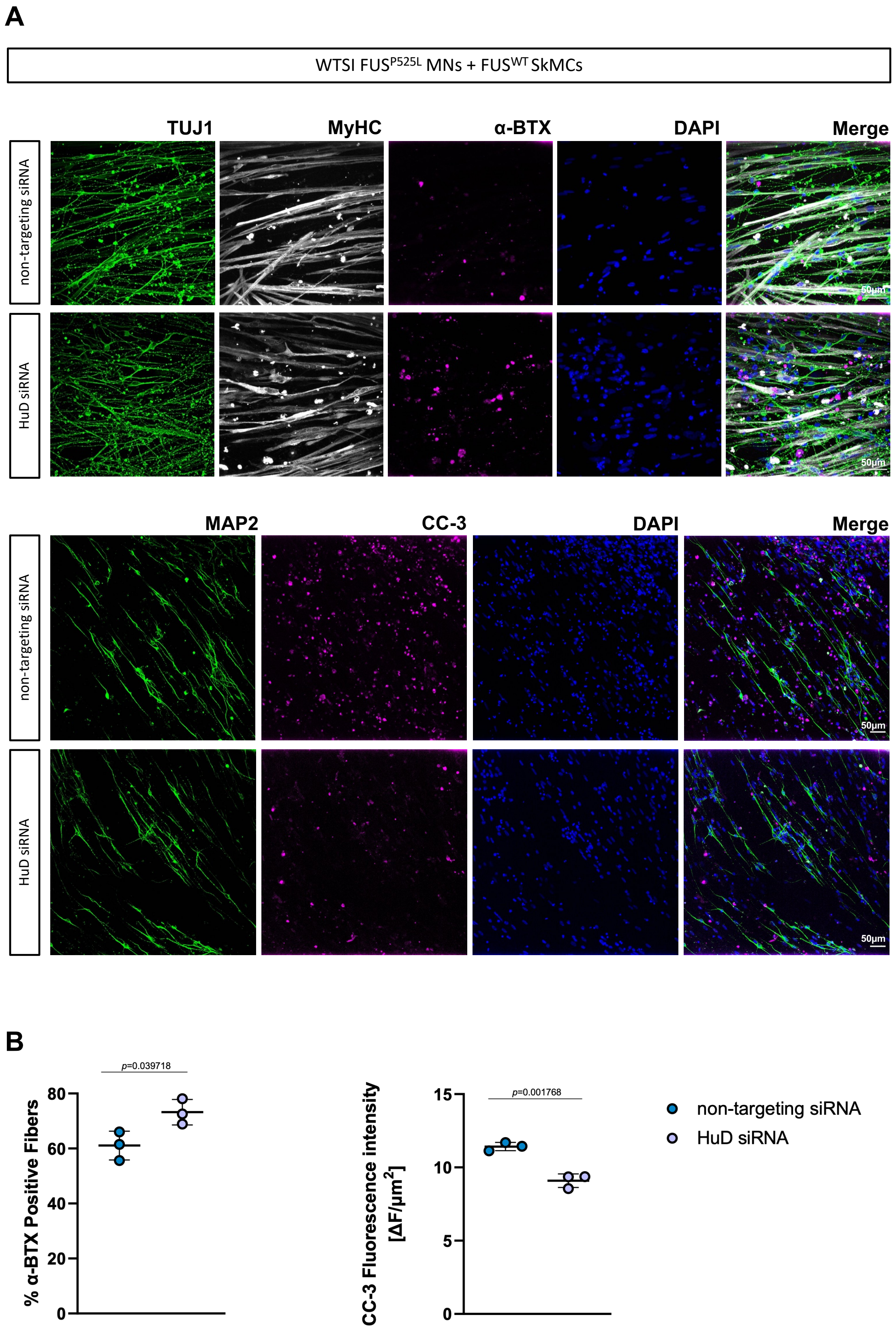


**Supplementary Figure S7. HuD knockdown in co-cultures containing WTSI-FUS$^{P525L}$ MNs. Related to Figure 5.**

(A) Representative images of day 14 co-cultures obtained with $FUS^{WT}$ SkMCs and WTSI-$FUS^{P525L}$ MNs transfected with non-targeting control or HuD siRNAs and stained with the indicated antibodies. Scale bar: 50µm. (B) The graphs report quantitative analysis of the percentage of α-BTX positive fibers (left) and CC-3 fluorescence intensity (right), at day 14 of co-culture as in (A). Each dot represents an individual batch of differentiated iPSCs transfected with the indicated siRNAs. For each replicate, 5-7 fields have been acquired for α-BTX positive fibers analysis and 8-10 fields for CC-3 immunofluorescence. The dot shows the average value. Error bars indicate standard deviation calculated on the average value of the replicates. Student's t-test; unpaired; two tails.

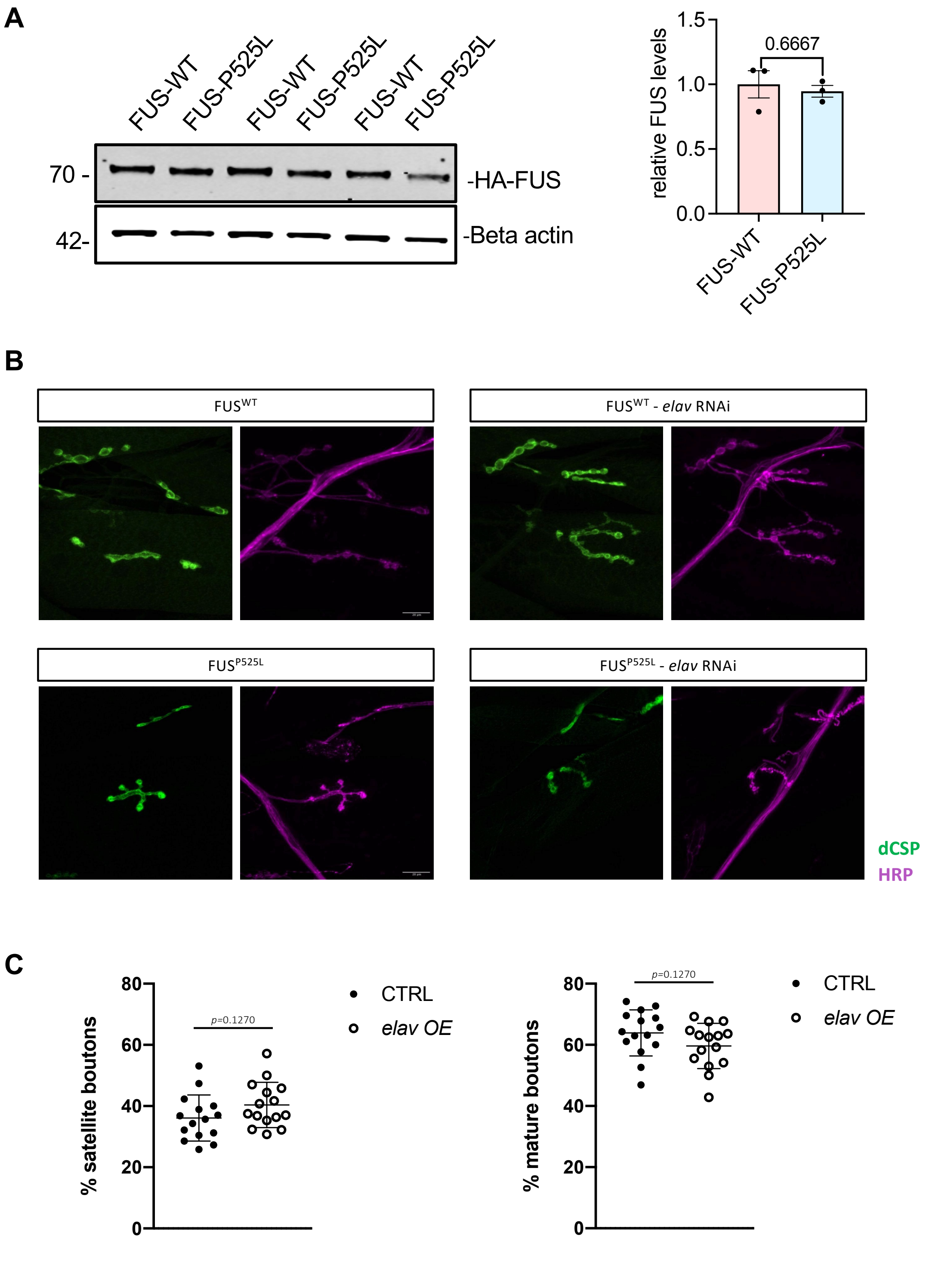


**Supplementary Figure S8. Expression of FUS proteins in *D. melanogaster* and NMJ analysis.** (A) Western blot of FUS protein level in adult fly heads (n = 3). Uncropped image is shown in Supplementary Figure S11. The graph on the right shows the quantification of western blots using ImageJ. The FUS expression was under a pan neuronal driver called the ELAV-Gene Switch. The F1 adult progenies were collected and fed with 5mM of RU-486 food for 1-6 days before performing the western blot. (B) Single panels of the IHC images shown in Figure 6. (C) The graphs show quantitative analysis of the percentage of satellite and mature boutons in larvae overexpressing *elav*

(*elav* OE) or Luciferase (LUC OE) as a control, respectively. Error bars indicate the mean and standard deviation, n = 6-10 larvae per genotype. Student's t-test, unpaired, two tails.

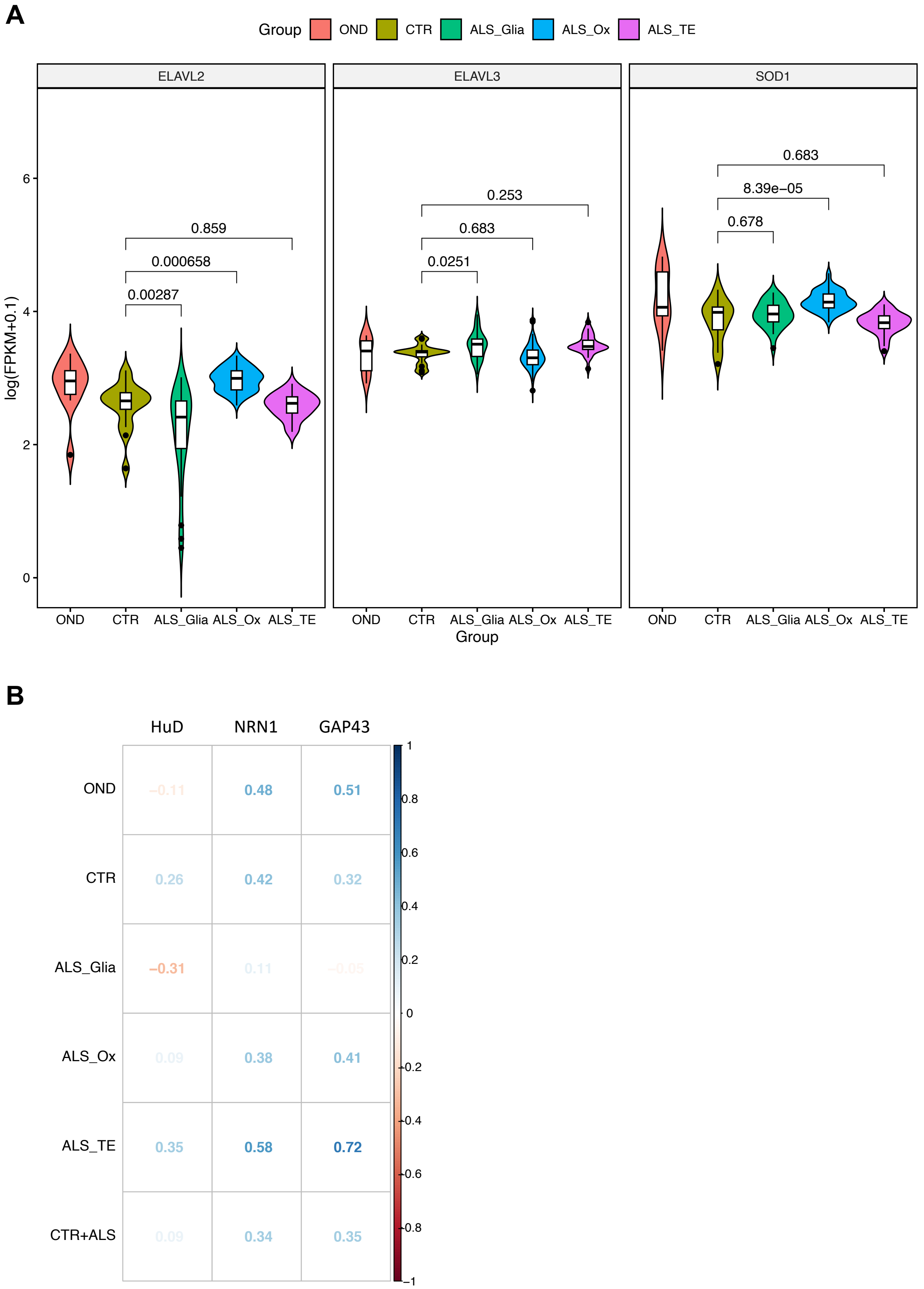


**Supplementary Figure S9. Analysis of nELAVLs and SOD1 expression in sporadic ALS patients and *HuD* variants. Related to Figure 7.**

(A) Violin plots showing the expression levels, reported as log-transformed FPKM values, of *ELAVL2* (*HuB*), *ELAVL3* (*HuC*) and *SOD1* in post-mortem sporadic ALS patients' cortex samples from (Tam et al., 2019). OND: other neurodegenerative diseases; CTR: healthy individuals; ALS_Glia: sporadic ALS patients with a signature of glial activation; ALS_Ox: sporadic ALS patients with a signature of oxidative and proteotoxic stress; ALS_TE: sporadic ALS patients with a signature of high retrotransposon expression. The adjusted *p*-values obtained from differential expression analyses are shown. (B) Correlation matrix reporting the Spearman's correlation coefficients calculated between the expression levels (FPKM) of *FUS* and those of *HuD*, *NRN1* and *GAP43* in samples from the NYGC consortium. The correlation values were calculated across various groups: OND, CTR, ALS_Glia, ALS_Ox, ALS_TE, and CTR+ALS (including CTR and all ALS patients' groups).

**A**

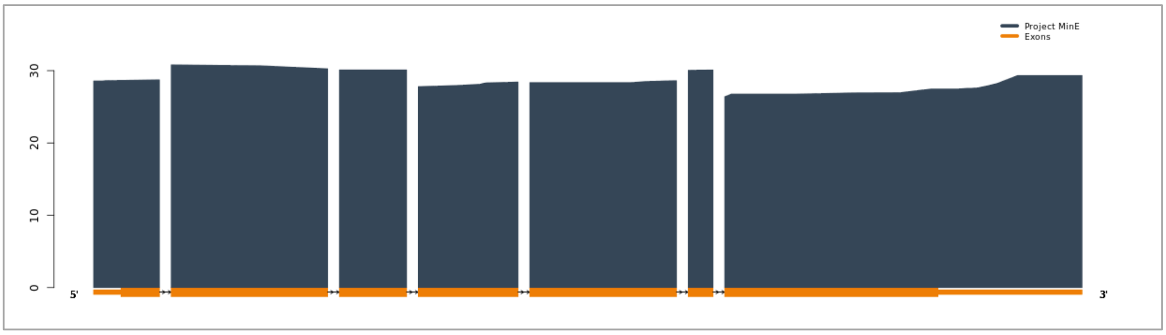


**B**

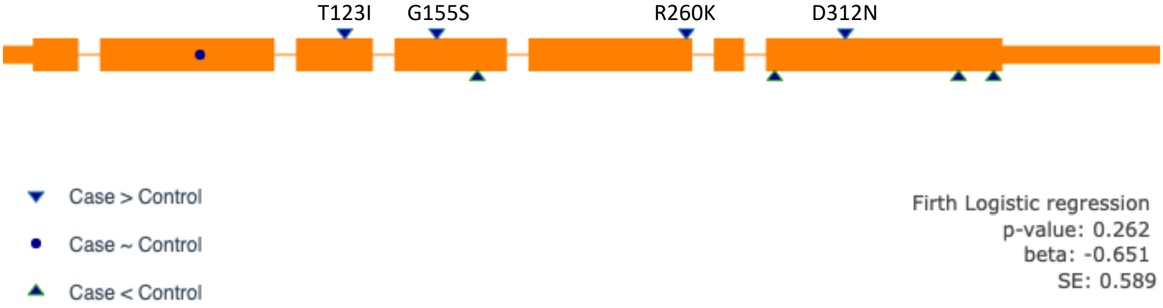


**Supplementary Figure S10. Variants in the HuD gene from Project MinE. Related to Figure 7.** Snapshots of the Project MinE data browser (http://databrowser.projectmine.com): coverage of the *HuD* gene (A) and exons (orange blocks) with the variants (triangles) that are reported in the current dataset (B). Variants that have been found with a higher frequency in ALS cases than controls are indicated.

A

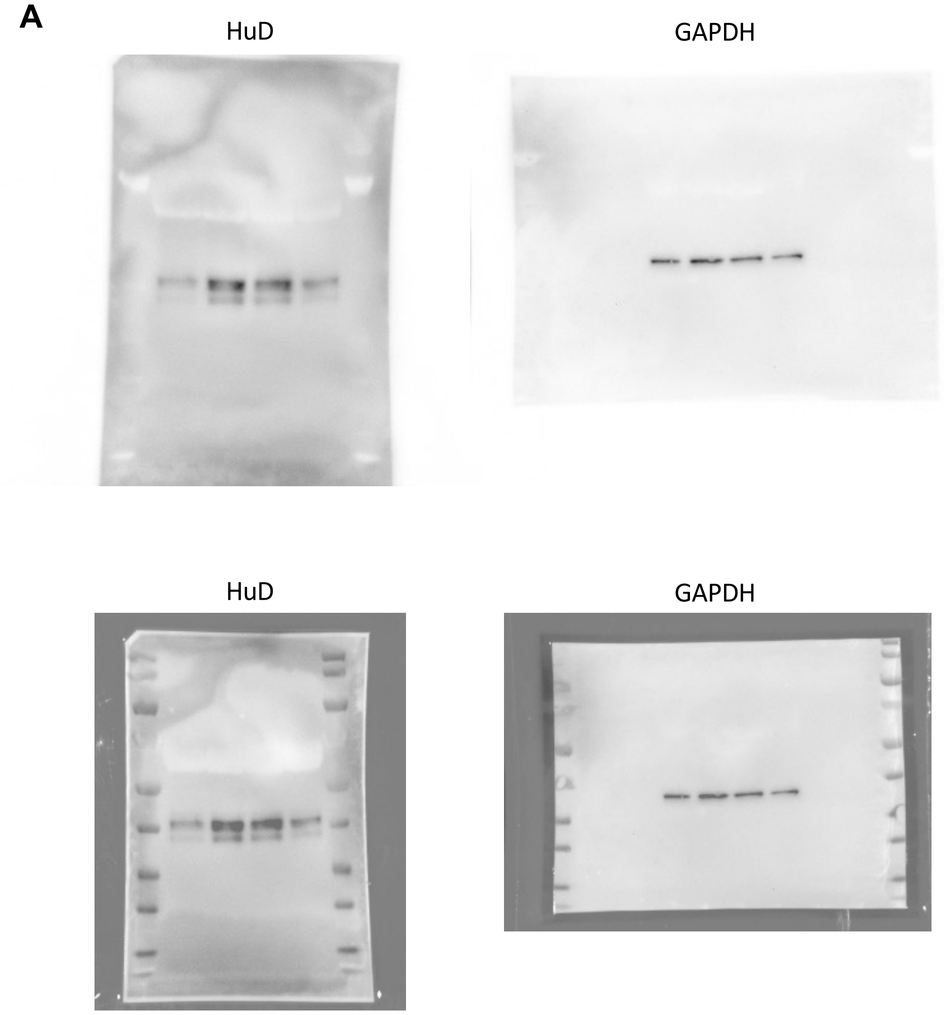


B

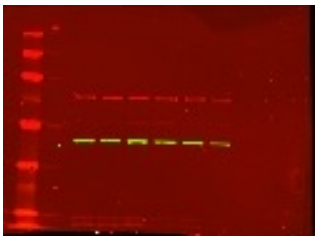

**Supplementary Figure S11. Uncropped images of the western blots and methods.**
(A) Images relative to Supplementary Figure S6 showing the overlap of the antibody signal and the molecular weight marker. An equal amount of protein samples was loaded in two gels, which were run and blotted in parallel. Filters were incubated with the following antibodies: anti-HuD (1:1000, sc-48421, Santa Cruz Biotechnology), anti-GAPDH (1:10000, sc-32233, Santa Cruz Biotechnology) primary antibodies and goat anti-mouse IgG (H+L) secondary antibody (32430, 1:10000, Thermo Fisher Scientific), as previously described (Garone et al., 2020). (B) Image relative to Supplementary Figure S8A. Fly heads were collected from each genetic cross and snap-frozen on dry ice. Western Blots were run in triplicate using biological replicates. Heads were crushed on dry ice and lysed with RIPA. Lysates were sonicated and centrifuged to remove debris. Supernatants were boiled in Laemmli Buffer for 5 minutes and loaded onto 4-12% NuPage Bis-Tris gels. The gels were transferred onto the nitrocellulose membrane using the iBlot2. The blots were blocked with milk

solution and further incubated in primary antibodies overnight. Primary antibodies used were rabbit beta-actin (1:1000; Cell Signaling Technology) and mouse anti-HA (1:1000; Sigma-Aldrich). Blots were washed and incubated at RT for one hour with anti-mouse IRDye 680D (1:10000) and anti-rabbit DYLight 800 (1:10000), and imaged on a Licor imager. Protein levels were quantified using NIH Image J software. Statistical analysis was performed with GraphPad Prism 8 software.

**Supplementary Table S1. List of primers used in this study.**

| Name | Sequence or assay |
|---|---|
| NANOG FW | CCAAATTCTCCTGCACGTGAC |
| NANOG RV | CACGTGGTTTCCAAACAAGAAA |
| ATP5O FW | ACTCGGGTTTGACCTACAGC |
| ATP5O RV | GGTACTGAAGCATCGCACCT |
| HuD FW | CAACCCCAGCCAGAAGTCCA |
| HuD RV | AGCCTGAACCTCTGAGCCTG |
| NRN1 FW | GGCTTTTCGGACTGTTTGCTCA |
| NRN1 RV | ATCCTCCCAGTATGTGCACACG |
| GAP43 FW | GAGGAGCCTAAACAAGCCGATG |
| GAP43 RV | GGGCACTTTCCTTAGGTTTGGT |
| MYH1 | Hs_MYH1_2_SG QuantiTect Primer Assay, QT01671005 |
| MYOG | Hs_MYOG1_SG QuantiTect Primer Assay, QT00001722 |
| DMD | Hs_DMD_1_SG QuantiTect Primer Assay, QT00085778 |
| OCT4 FW | ATGCATTCAAACTGAGGTGCCTGC |
| OCT4 RV | AACTTCACCTTCCCTCCAACCAGT |
| PAX7 FW | CAGAGGACCAAGCTGACAGA |
| PAX7 RV | CTGGCAGAAGGTGGTTGAAC |
| SOD1 FW | TGAAGGTGTGGGGAAGCATT |
| SOD1 RV | TCTCTTCATCCTTTGGCCCA |
| TUBB3 FW | CCCGGAACCATGGACAGTGT |
| TUBB3 RV | TGACCCTTGGCCCAGTTGTT |
